# Direct sensitizing and activating effects of interleukin 31 are restricted to a single, functionally and transcriptionally classified porcine DRG neuron subtype

**DOI:** 10.64898/2026.04.10.717660

**Authors:** Zeinab Abbasi, Marc Behrendt, Sabrina da Silva Soares, Roman Rukwied, Martin Schmelz, Hans Jürgen Solinski

## Abstract

Interleukin-31 (IL-31) drives chronic pruritus in patients with dermatological and even certain systemic diseases. However, fast-onset anti-pruritic effects of blocking IL-31 receptors, for instance with nemolizumab in atopic dermatitis patients, are incompletely understood, in part due to ethical restrictions in humans and species differences to mice. Therefore, we used sensory neuron cultures from pig to investigate direct neuronal IL-31 effects. We first mapped functional characteristics of afferents encoding histamine itch in humans onto a recently established transcriptome-based DRG neuron taxonomy to identify pig pruriceptors. IL-31 acutely sensitized responses to repeated pruritogen and electrical stimulation only in these histamine- and capsaicin-responsive pruriceptors and also activated these afferents with silent nociceptor phenotype *in vivo* as validated by dermal axon-reflex erythema measurements. Thus, our data functionally and transcriptionally identifies the likely sensory neuron class underlying IL-31-driven chronic pruritus and opens a perspective for translational research on distinct neuronal classes differentially driving skin inflammation and clinical chronic pruritus via specific neuro-immune signaling patterns.

## Introduction

Interleukin 31 (IL-31) is considered one of the key molecular drivers of chronic pruritus (1). IL-31 levels in plasma and skin correlate well with chronic pruritus intensity in dermatological and even certain systemic diseases (2–5). In the clinics, inhibition of IL-31 signaling through blockage of its heterodimeric receptor IL31Rα/OSMRβ with nemolizumab (blocking IL31Rα) or vixarelimab (blocking OSMRβ) results in fast-onset reduction of itch before ameliorating effects on skin inflammation are detected (6–8). These fast drug effects on chronic pruritus suggest inhibition of ongoing IL-31 signaling in somatosensory neurons as one therapeutic mechanism. Accordingly, both IL-31 receptor subunits have been detected in murine transient receptor potential vanilloid subfamily member 1 expressing (TRPV1^+^) neurons and this expression pattern was likewise established in other species, including humans (9–12). However, the exact neuronal mechanisms by which IL-31 promotes chronic or even acute pruritus are only incompletely understood, in part due to ethically restricted access to human somatosensory neurons. For instance, intradermal IL-31 delivery has been reported to induce immediate site-directed scratching in mice (12), indicative for a role as a *bona fide* pruritogen. But other reports in mice, dogs, monkeys and humans point to the lack of such an immediate response and rather reported site-directed but delayed scratching or an anatomically generalized scratching response, potentially involving neuronal sensitization (13–19).

In primates and humans, at least three types of skin afferent nerve fibers have been implicated in acute pruritus (20–22). Overall, afferent nerve fiber types in humans are distinguished based on functional properties of their sensory ending and axon, including mechano- and heat-sensitivity, conduction velocity, thresholds and activation patterns to fast rectangular or slow sinusoidal electrical stimulation and following frequency to fast rectangular electrical pulses (23–30). Functional features of afferents encoding histaminergic itch include the lack of mechano-sensitivity, low conduction velocity, broad action potentials and, importantly, high rectangular but low sinusoidal electrical thresholds (20, 26). However, it is currently unclear how functionally defined afferent fiber types with or without a role in pruritus relate to dorsal root ganglion neuron types defined structurally by post-mortem single-cell transcriptomics (11, 31–33). Yet, linkage of transcriptional and functional information on sensory neurons would be highly valuable in order to devise new treatment strategies for chronic itch patients.

Herein, we propose the back-translation of tests used to functionally categorize human skin afferents to a non-human model system that allows a simultaneous measurement of chemical response patterns, linking them to transcriptionally defined somatosensory neuron types. We chose domestic pigs for this purpose due to the established similarities in skin physiology (34, 35), inflammatory skin pathophysiology (36, 37) and, importantly, skin afferent neurobiology (24, 28, 38), the latter resulting in a similar portfolio of functionally defined skin afferent nerve fiber types as in humans. We use this approach to investigate neuronal structure-function relations using acute histamine responses as functional marker and subsequently study direct effects of IL-31 in transcriptionally defined porcine sensory neurons to identify those most relevant for IL-31-driven chronic pruritus.

## Results

### Chemical response patterns as proxy for transcriptionally defined neuron-types

We took advantage of a recently published single-nucleus RNA sequencing (snRNAseq) dataset of the porcine DRG (13) and investigated the expression of chemically targetable receptors within transcriptionally defined DRG neuron clusters. Based on the knowledge about chemical responses in fibers encoding histamine itch in humans, we focused our investigation on the genes *HRH1* and *TRPV1*. As shown in figure 1A, *HRH1* expression is largely confined to a single transcriptionally defined neuron type that was annotated as C-OSMR-SST and has high transcriptional similarity to *SST* expressing DRG neurons in other species, including humans (13). *TRPV1* expression was expectedly detected in multiple C-fibers and one putative Aδ-fiber cluster but was virtually absent in all Aβ-fiber and some Aδ/C-fiber clusters. Importantly, C-OSMR-SST neurons with high *HRH1* expression also co-expressed *TRPV1*. We therefore hypothesized that we could use a consecutive stimulation with histamine and capsaicin to group porcine DRG neurons based on their expression of *HRH1* and *TRPV1*. We expected that histamine responsive neurons would show a high degree of capsaicin responsiveness corresponding to the *HRH1*/*TRPV1* co-expression in C-OSMR-SST neurons. In addition, we expected to find large numbers of histamine insensitive neurons that we could subdivide based on their capsaicin response. These histamine-insensitive groups of neurons would represent mixtures of several transcriptionally defined neuron groups with capsaicin positive nociceptors consisting primarily of several C-TAC1 and the Aδ-TRPV1 clusters and capsaicin non-responders consisting of the remaining C/Aδ- and all Aβ-clusters. Indeed, Fluo8-based calcium imaging of primary porcine DRG neuron cultures, performed on the following day after dissociation, (Fig. 1B-D) segregated neurons into three groups with the expected response patterns to consecutive stimulation with histamine (200 µM) and capsaicin (10 µM) and we will refer to these three groups throughout this paper as CAP^+^/His^+^, CAP^+^ and CAP^−^ neurons. In preparations from 12 pigs, we measured a total of 589 neurons. 70 neurons (11.9%) responded to histamine and >87% of these neurons also responded to capsaicin. Capsaicin activated 258 neurons (43.8%) but only 24.4% of these neurons also responded to histamine, collectively indicating that histamine responsiveness is largely confined to a subpopulation of CAP^+^ neurons. 324 neurons (55.0%) did neither respond to histamine or capsaicin. Of note, in order to functionally also detect neurons expressing low levels of *TRPV1,* we used a high concentration of capsaicin. We anticipated that this would likely produce acute excitotoxicity and indeed found that a subsequent response to a high KCl solution (+55 mM) was prohibited in 84.6% of all capsaicin responding neurons.

**Figure 1.**
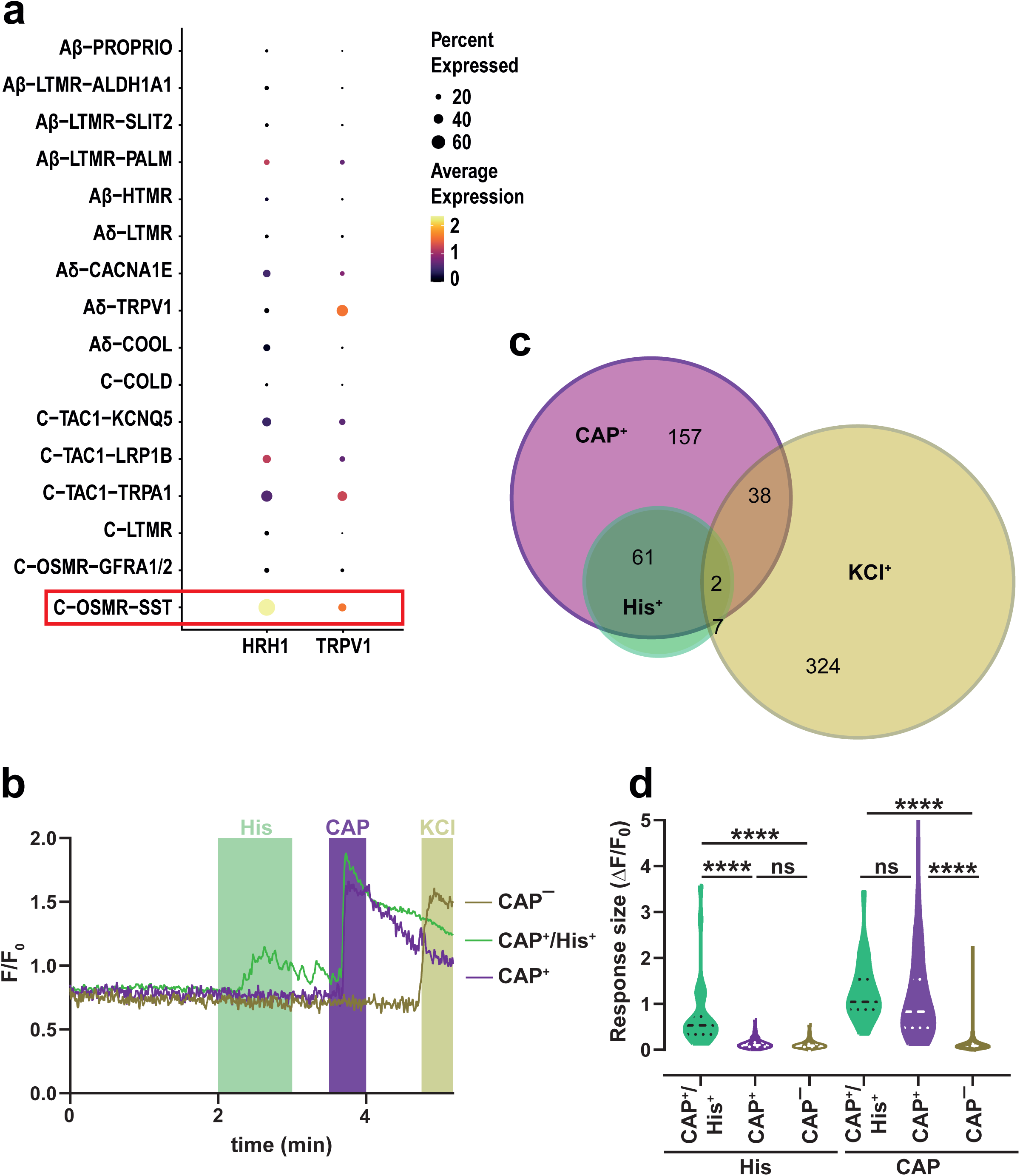
Chemical response pattern-based classification of porcine DRG neurons. a) Expression of *HRH1* and *TRPV1* in porcine DRG neuron types, as assessed by reanalysis of snRNAseq data (13) of porcine thoracic and lumbar DRGs, shown by a dot plot. C-OSMR-SST neurons co-expressing high levels of *HRH1* and *TRPV1* are highlighted with a red rectangle. b) Calcium traces of three single neurons, representative for the investigated neuron groups. Chemical stimulations are indicated with shaded areas, histamine (His, 200 µM): green, capsaicin (CAP, 10 µM): purple, KCl (+55 mM): olive. c) Venn diagram with neuron numbers indicating population sizes of the observed response patterns (n = 12 pigs, 589 cells). d) Net response sizes to chemical stimulation with histamine and capsaicin in all three neuron groups are depicted as violin plot (dashed line: median, dotted lines: quartiles), Kruskal-Wallis ANOVA with Dunn’s post-hoc test.

However, we deem this phenomenon as acceptable, as we are applying capsaicin and KCl only at the end of each protocol and as we do not use the KCl response in capsaicin responding neurons for cell filtering or grouping. In order to test the stringency of neuron grouping based on chemical response patterns (see methods), we compared the net response sizes of calcium signals to histamine and capsaicin superfusion among our three neuron groups (Fig. 1D). As expected, histamine responses were significantly higher in CAP^+^/His^+^ neurons in comparison to both other neuron groups (p < 0.0001, Kruskal-Wallis + Dunn’s post-hoc test), while capsaicin responses were significantly higher in CAP^+^/His^+^ and CAP^+^ neurons in comparison to CAP^−^ neurons (p < 0.0001, Kruskal-Wallis + Dunn’s post-hoc test) but indistinguishable between themselves (p > 0.9999). This indicated that our method resulted in robust cell groups, representing transcriptionally defined porcine DRG neuron types/groups.

### Differential responses to rectangular electrical stimulation

Next, we assessed responsiveness to extracellular electrical stimulation of porcine DRG neuron groups. We first applied extracellular electrical stimuli using the classic rectangular stimulation wave form (2 ms pulse width) at very high intensity (100 mA) to induce a defined number of 1, 2 or 4 action potentials (APs, Fig. 2A upper panel). This stimulation allowed us to measure calcium transients associated with a single AP and, in addition, how such signals superimpose if 2-4 APs are triggered at 4 Hz frequency. As shown in figure 2A, this stimulation regime allowed to trigger repetitive AP-associated calcium transients in all three neuron groups. On the group level, the number of responding neurons increased with the number of stimulation pulses (Fig. 2B; CAP^+^/His^+^: 25-30 out of 43 cells, CAP^+^: 61-85 out of 123 cells, CAP^−^: 105-183 out of 234 cells), indicating a higher detection sensitivity when multiple AP-associated calcium transients superimpose. Although we noticed this behavior for all neuron groups, proportions of responding neurons only increased significantly within CAP^+^ and CAP^−^ neurons (in CAP^+^/His^+^, CAP^+^, CAP^−^ neurons: χ2 = 1.255, 9.666, 55.16; p = 0.2626, 0.0019, <0.0001; chi-square test for trend). However, when focusing only on stimulation with a single pulse to remove potentially confounding effects of differing following frequencies at stimulations with multiple stimuli, we did not detect a systematic difference in the population sizes of responding neurons between the three neuron groups (χ2 = 2.805, p = 0.2460, chi-square test). Next, we considered only neurons responding to all three rectangular stimulation pulses (see methods; n = 24, 59, 97 neurons in CAP^+^/His^+^, CAP^+^, CAP^−^ neurons) and extracted various characteristics of the measured electrically evoked calcium transients to allow quantitative comparisons between the neuron groups. We focused on the stimulation-induced F/F_0_ increases (Fig. 2C) as a proxy for the response size and on the time for a signal to decline to 50% (T50) of these increases (Fig. 2D) as a proxy for the response duration. After a single stimulation pulse (Fig. 2C, left panel), the resulting AP-associated calcium response sizes (CAP^+^/His^+^: 0.29 ± 0.04, CAP^+^: 0.28 ± 0.02, CAP^−^: 0.24 ± 0.01) did not differ significantly between the three neuron groups (H (2) = 2.96, p = 0.2274, Kruskal-Wallis). When we compared those sizes for multiple stimulation pulses (Fig. 2C, middle and right panel), a significant difference emerged (2 pulses: H (2) = 10.36, p = 0.0056; 4 pulses: H (2) = 15.23, p = 0.0005; Kruskal-Wallis). These differences were solely driven by lower calcium response sizes in CAP^−^ neurons (-26% for 2 pulses, -34% for 4 pulses, both vs. average of CAP^+^/His^+^ and CAP^+^ neurons) that also reached statistical significance in pairwise comparisons to CAP^+^ neurons (CAP^−^ vs CAP^+^/His^+^ p = 0.1353/0.0474 and CAP^−^ vs CAP^+^ p = 0.0089/0.0009, each for 2/4 pulses, Dunn’s post-hoc test). Since the response sizes to a single stimulation pulse did not differ among neuron groups and since this result could be reproduced by normalizing responses sizes on the single-cell level to calcium transients associated with a single stimulation pulse (Fig. S1), the most likely explanation for this result is a lesser degree of calcium transient superimposition in CAP^−^ neurons. Functionally, such behavior fits to Aβ-fibers, known to transmit AP trains with frequencies up to 400 Hz (25), that are only contained in CAP^−^ neurons. Unexpectedly, T50 duration differentiated between the neuron classes, segregating CAP^+^/His^+^ neurons from the two other neuron types (Fig. 2D). For 1, 2 and 4 stimulation pulses, T50 of CAP^+^/His^+^ neurons was the longest among the three neuron groups (+66/65/45% for 1/2/4 pulses, all vs. average of CAP^+^ and CAP^−^ neurons) and this difference was overall statistically significant in each stimulation regime (p = < 0.0001 for 1/2/4 pulses, Kruskal-Wallis) and persisted in pairwise comparisons to CAP^−^ (p = < 0.0001 for 1/2/4 pulses, Dunn’s post-hoc test) and CAP^+^ (p = 0.0004/0.0027/0.0037, Dunn’s post-hoc test) neurons. In addition, we also detected a trend for longer T50 duration in CAP^+^ neurons when comparing them with CAP^−^ neurons but this difference depended on the number of stimulation pulses and only reached statistical significance for 4 stimuli (>0.9999/0.2257/0.0136, Dunn’s post-hoc test).

**Figure 2.**
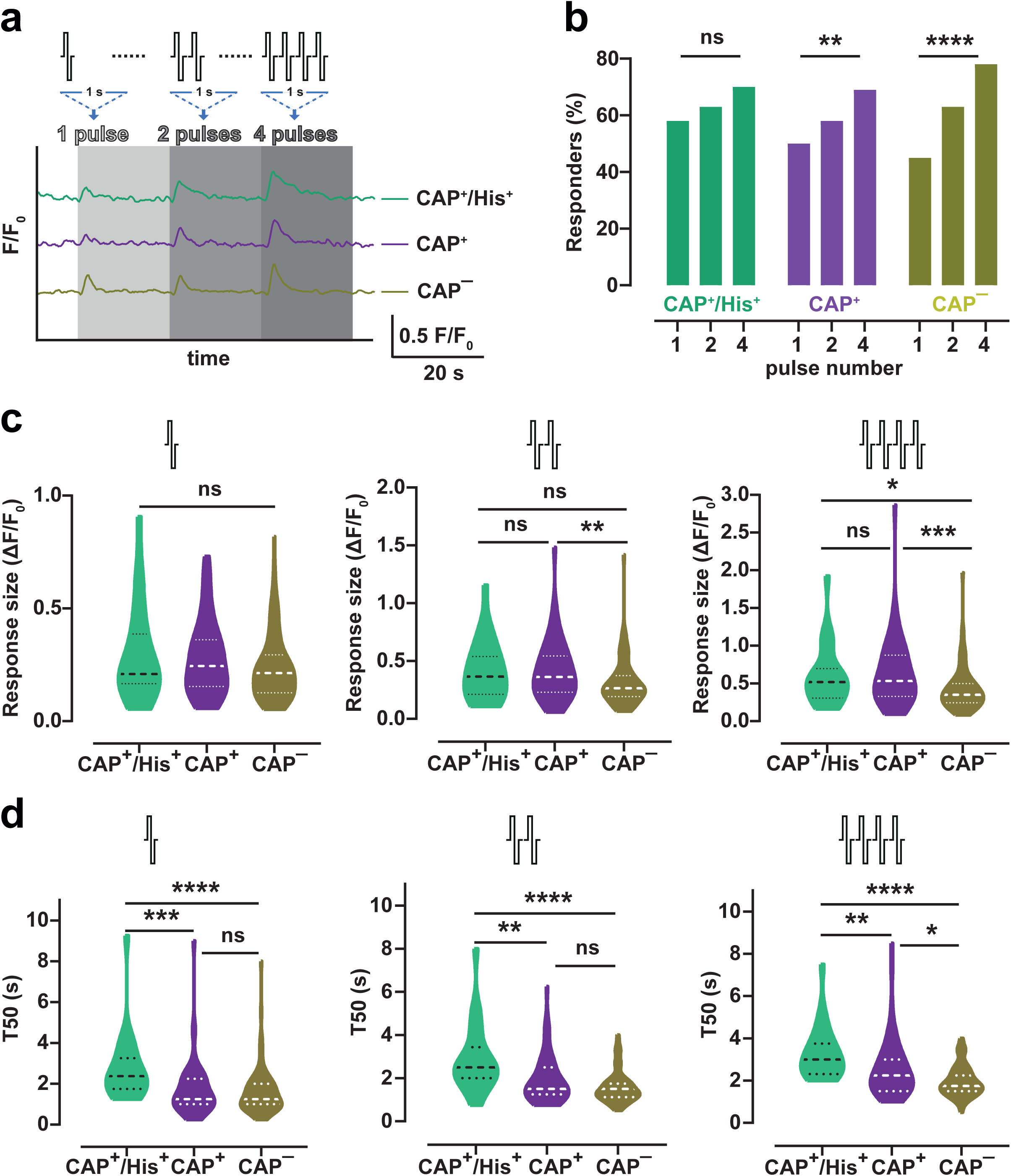
AP-associated calcium transients differ between porcine DRG neuron types. a) Rectangular electrical stimulation (100 mA) applied in 1, 2, and 4 pulse patterns induces action potential-associated calcium transients in single neurons representative for the three neuron groups (n = 9 pigs). b) Proportion of neurons responding to 1, 2, and 4 pulses of rectangular electrical stimulation (total neuron number: n = 43 (CAP^+^/His^+^), n = 123 (CAP^+^), and n = 234 (CAP^−^)), chi-square test for trend. c) Maximum size of calcium responses to rectangular electrical stimulation with 1 (left), 2 (middle) and 4 (right) stimuli. Data for cells responding to all 3 stimulation patterns (n = 24 (CAP^+^/His^+^), n = 59 (CAP^+^), and n = 97 (CAP^−^)) is depicted as violin plot (dashed line: median, dotted lines: quartiles), Kruskal-Wallis ANOVA with Dunn’s post-hoc test. d) Half-decay time (T50) of calcium responses to rectangular electrical stimulation with 1 (left), 2 (middle) and 4 (right) stimuli. Data for same cells plotted in C is depicted as violin plot (dashed line: median, dotted lines: quartiles), Kruskal-Wallis ANOVA with Dunn’s post-hoc test.

### Differential responses to sinusoidal electrical stimulation

Given that mechano-insensitive C-chemonociceptors show a strong preference for slow depolarizing electrical stimuli with sinusoidal stimulation wave forms (250 ms duration, 4 Hz, slow sine) *in vivo* (23), we next stimulated porcine DRG neurons with extracellular electrical stimuli that follow such a sine wave shape (Fig. 3A). Based on frequency-response relationships measured in human and pig skin (29, 30), we also applied a single complete sine wave cycle at 20 ms duration (50 Hz, fast sine) to stimulate C-fiber neurons most efficiently. Beginning with a fast sine wave cycle, these two paradigms were delivered alternatingly at increasing intensities, starting with 5 mA and increasing in 5 mA steps up to 50 mA with 20 second intervals (specimen responses in Fig. 3B). First, we compared responses between neuron groups within each of the two sinusoidal wave forms. To keep with our analysis pipeline employed for rectangular electrical stimulation, we extracted net response amplitudes (F/F_0_ increases) and signal width (T50) at maximum stimulation intensity (50 mA) considering only responding neurons (see methods, n = 34/15, 73/51, 162/128 neurons for slow/fast sine in CAP^+^/His^+^, CAP^+^, CAP^−^ neurons). Overall, at this stimulation intensity the results of sinusoidal electrical stimulation mimicked the results of rectangular electrical stimulation, displaying similar response amplitudes but prolonged response times in CAP^+^/His^+^ neurons. In detail, we found that response amplitudes were similar between the three neuron groups for both sine waves (Fig. 3C; fast sine: H (2) = 0.5019, p = 0.7781; slow sine: H (2) = 5.733, p = 0.0569; both Kruskal-Wallis). The signal width for sinusoidal stimulation-induced calcium transients was substantially longer in CAP^+^/His^+^ neurons (Fig. 3D; fast sine: +43%, slow sine: +57%, both vs average of CAP^+^ and CAP^−^ neurons), which reached statistical significance overall (fast sine: H (2) = 7.525, p = 0.0232; fast sine: H (2) = 22.58, p < 0.0001; Kruskal-Wallis) and in both pairwise comparisons (vs CAP^+^ p = 0.0395/< 0.0001, vs CAP^−^ p = 0.0212/< 0.0001 for fast/slow sine; Dunn’s post-hoc test). This finding substantiates that electrical stimulation-induced calcium transients in CAP^+^/His^+^ neurons are generally longer lasting, irrespective of the used stimulation wave form. Second, we compared both sinusoidal wave forms within each of the three neuron groups. For this comparison, we considered neurons as responding if they showed a significant calcium transient to any of the 10 stimulation intensities. For all neuron groups, the percentage of responding neurons was larger when stimulated with the slow sine wave compared to the fast sine wave (Fig. 3E). However, CAP^+^/His^+^ neurons displayed the largest percentage of responding neurons to slow and inversely the lowest percentage of responding neurons to fast sine wave stimulation, resulting in a significantly larger increase in responder incidence for slow vs fast sine waves of + 48% (CAP^+^/His^+^: p < 0.0001, CAP^+^: p = 0.2424, CAP^−^: p = 0.1008; Fisher’s exact test) and suggesting that slower sine waves are better suited to activate CAP^+^/His^+^ neurons. Therefore, we finally calculated the slow sine wave preference (see methods) as an aggregate measure of thresholds to both sine waves for each individual neuron. This analysis showed that for CAP^+^/His^+^ neurons, thresholds to the slow sine wave were indeed much lower (on average -15 ± 3.5 mA) as compared to CAP^+^ (-4 ± 2.0 mA) and CAP^−^ neurons (-1 ± 1.3 mA) and this striking difference was highly significant (Fig. 3F; H (2) = 17.04, p = 0.0002, Kruskal-Wallis; CAP^+^/His^+^ vs CAP^+^: p = 0.0125, CAP^+^/His^+^ vs CAP^−^: p = 0.0001, both Dunn’s post-hoc test). This strongly suggests that functional phenotypes present in thin skin afferent nerve endings *in vivo* can also be found in neuronal cell bodies *in vitro.* Thus, the characteristics of CAP^+^/His^+^ neurons encompass both, the transcriptional pattern of C-OSMR-SST neurons and the functional pattern of mechanically insensitive histamine itch encoding C-chemonociceptors.

**Figure 3.**
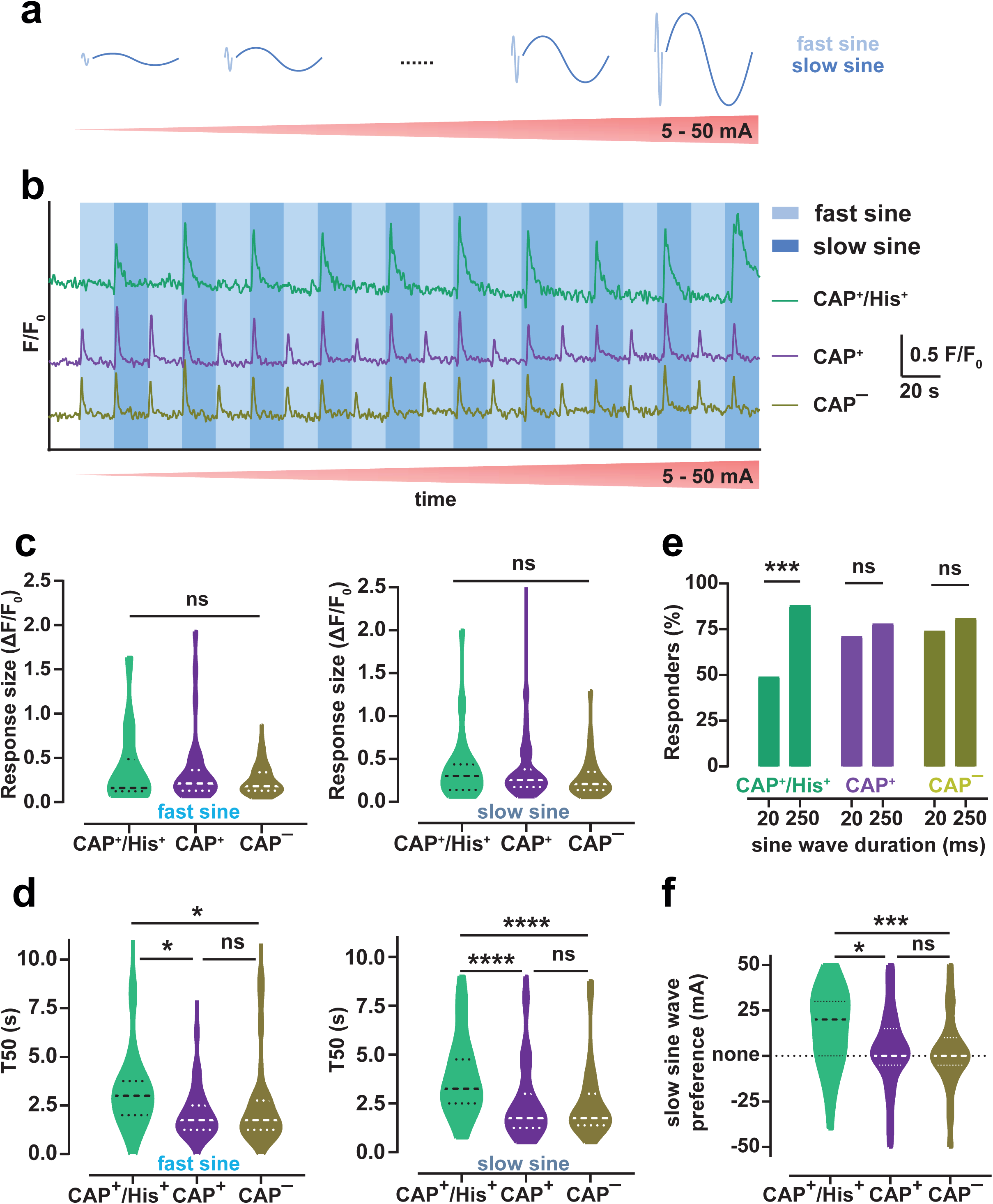
Calcium transients elicited by sinusoidal electrical stimulation differ between porcine DRG neuron types. a) Sinusoidal electrical stimulation protocol with intensities varying in 5 mA steps from 5 to 50 mA and alternating between fast (20 ms) and slow (150 ms) sine waves at each intensity level is depicted (n = 9 pigs). b) Calcium traces in response to sinusoidal electrical stimulation in single neurons representative for the three neuron types, fast sine waves shaded by light blue, and slow sine waves shaded by dark blue. Note the strong preference for slow sine wave stimulation in the CAP^+^/His^+^ neuron not visible in neurons from both other groups. c) Maximum size of calcium responses to maximum current intensity (50 mA) electrical stimulation with fast (left) and slow (right) sine waves. Data for cells responding to fast (n = 15 (CAP^+^/His^+^), n = 40 (CAP^+^), and n = 127 (CAP^−^)) or slow (n = 34 (CAP^+^/His^+^), n = 65 (CAP^+^), and n = 161 (CAP^−^)) sine wave stimulation is depicted as violin plot (dashed line: median, dotted lines: quartiles), Kruskal-Wallis ANOVA. d) Half decay time (T50) of calcium responses to maximum current intensity (50 mA) electrical stimulation with fast (left) and slow (right) sine waves. Data for same cells plotted in C is depicted as violin plot, Kruskal-Wallis ANOVA with Dunn’s post-hoc test. e) The proportion of neurons responding to any intensity of electrical stimulation with fast or slow sine waves (total neuron number: n = 43 (CAP^+^/His^+^), n = 123 (CAP^+^), and n = 240 (CAP^−^)), Fisher’s exact test. f) Slow sine wave preference (difference of fast minus slow sine wave thresholds, non-responder thresholds were set to 55 mA) was calculated for cells responding to any intensity of electrical stimulation with fast or slow sine waves (n = 39 (CAP^+^/His^+^), n = 107 (CAP^+^), and n = 216 (CAP^−^)). Data is given as violin plot (dashed line: median, dotted lines: quartiles), Kruskal-Wallis ANOVA with Dunn’s post-hoc test.

### Direct stimulation of porcine DRG neurons by IL-31

Direct and ongoing excitation of itch-encoding somatosensory neurons by IL-31 has been hypothesized to foster chronic pruritus. However, local skin stimulation with IL-31 in healthy individuals produces only weak and delayed pruritus in humans or delayed scratching behavior in large animals (14–17). We therefore directly investigated IL-31 signaling events in sensory neurons and tried to assign them to neuronal classes identifiable through functional and transcriptome-based classification.

A pre-requisite for any direct IL-31-induced effects in porcine DRG neurons is the expression of both subunits of the IL-31 receptor, IL31RA and OSMRβ (39), as well as the primary signal-transducing kinase JAK1. Pig snRNAseq data (13) suggest that C-OSMR-SST neurons are virtually the only transcriptionally defined neuron type expressing mRNA for these three proteins (Fig. 4A), making them prime neuronal candidates for IL-31-induced signaling. IL-31, IL31RA and OSMRβ show only a modest degree of evolutionary conservation, as measured by a comparison of the amino acid sequence of porcine, murine and human proteins (Table S1-3), suggesting a lack of cross-species activity of IL-31 and its receptor complex. We therefore obtained recombinant porcine IL-31 (rpIL-31, see methods) and tested its activity by measuring induction of the most canonical IL-31-induced signaling pathway, STAT3 activation (39), in a recombinant expression system (Fig. S2A). Of note, initial tests corroborated our suspicion of a lack of cross-species activity, as recombinant human IL-31 did not show any appreciable activity when stimulating recombinantly expressed porcine IL31RA and OSMRβ (Fig. S2A, p > 0.9999, Sidak’s post-hoc test), while rpIL-31 strongly stimulated reporter activity but only when the porcine IL-31 receptor complex was expressed (p < 0.0001, Sidak’s post-hoc test). Measuring concentration-response relationships of rpIL-31 at its cognate porcine receptor complex in this recombinant expression system, we estimate a potency of our rpIL-31 of 48 ± 5 ng/ml (Fig. S2B). Based on these results, we chose to conduct further experiments with porcine DRG neurons at a dose equivalent to an EC_75_ value (272 ng/ml) for STAT3 activation in the recombinant expression system. First, we studied whether acute stimulation of pig DRG neurons with rpIL-31 produces calcium transients. Overall, this was a rare but well detectable phenomenon, happening in ∼5% of the tested neurons (10 out of 202). Based on our single-nucleus expression analysis, we expected such responses to be restricted to CAP^+^/His^+^ neurons. And indeed, 26% of CAP^+^/His^+^ neurons displayed acute IL-31-induced calcium transients (Fig. 4B+C), while this happened in only 4.5% of CAP^+^ and was never detected in CAP^−^ neurons (Fig. 4C+D, χ2 = 26.74, p < 0.0001, chi-square test; CAP^+^/His^+^ vs CAP^+^: p = 0.018, CAP^+^/His^+^ vs CAP^−^: p < 0.001, CAP^+^ vs CAP^−^: p = 0.147, all Fisher’s exact test with p-values adjusted by Bonferroni’s method). This result indicated that rpIL-31 can directly stimulate C-OSMR-SST neurons. However, *IL31RA*, *OSMR*β and *JAK1* were overlappingly expressed in >80% of C-OSMR-SST neurons (Fig. 4A), suggesting that a much higher percentage of CAP^+^/His^+^ neurons could respond to rpIL-31 but, potentially, engaging other intracellular signaling pathways. Calcium mobilization in sensory neurons by rpIL-31 does not necessarily denote AP generation. We therefore tested next whether stimulation of C-OSMR-SST fibers in the pig skin *in vivo* produces neuronal activity. As indirect readout for chemonociceptor activity, we measured AP-dependent axon-reflex vasodilation in the skin (40) with a speckle laser. Because we detected rpIL-31-induced calcium transients only in 25% of CAP^+^/His^+^ neurons, histamine injections were used to estimate maximum axon-reflex erythema sizes and kinetics. Histamine injections substantially increased areas of the axon-reflex vasodilation in comparison to control injections with PBS for up to 4 minutes post injection (Fig. 4D+E, p < 0.0001, Sidak’s post-hoc test). Strikingly, intradermal rpIL-31 injections also produced an immediate and significant axon-reflex erythema (Fig. 4D+E), substantially larger than the injection bleb and the local erythema after control injections (p < 0.0001, Sidak’s post-hoc test). In good agreement with the different number of responsive neurons *in vitro*, histamine-induced axon-reflex erythema *in vivo* was indeed larger and exceeded the rpIL-31-induced effect 1.9-fold. Importantly, the size of the hyperperfused area increased with increasing rpIL-31 doses, following a sigmoidal dose-response relationship with an apparent potency of 5 ± 10 ng/ml (Fig. 4F). Because the axon-reflex in human (40) and pig skin (41) is an *in vivo* hallmark of mechano-insensitive C-chemonociceptors, we take this result as orthogonal functional evidence for C-OSMR-SST neurons being the transcriptionally defined correlate of mechano-insensitive histamine itch-encoding skin afferents.

**Figure 4.**
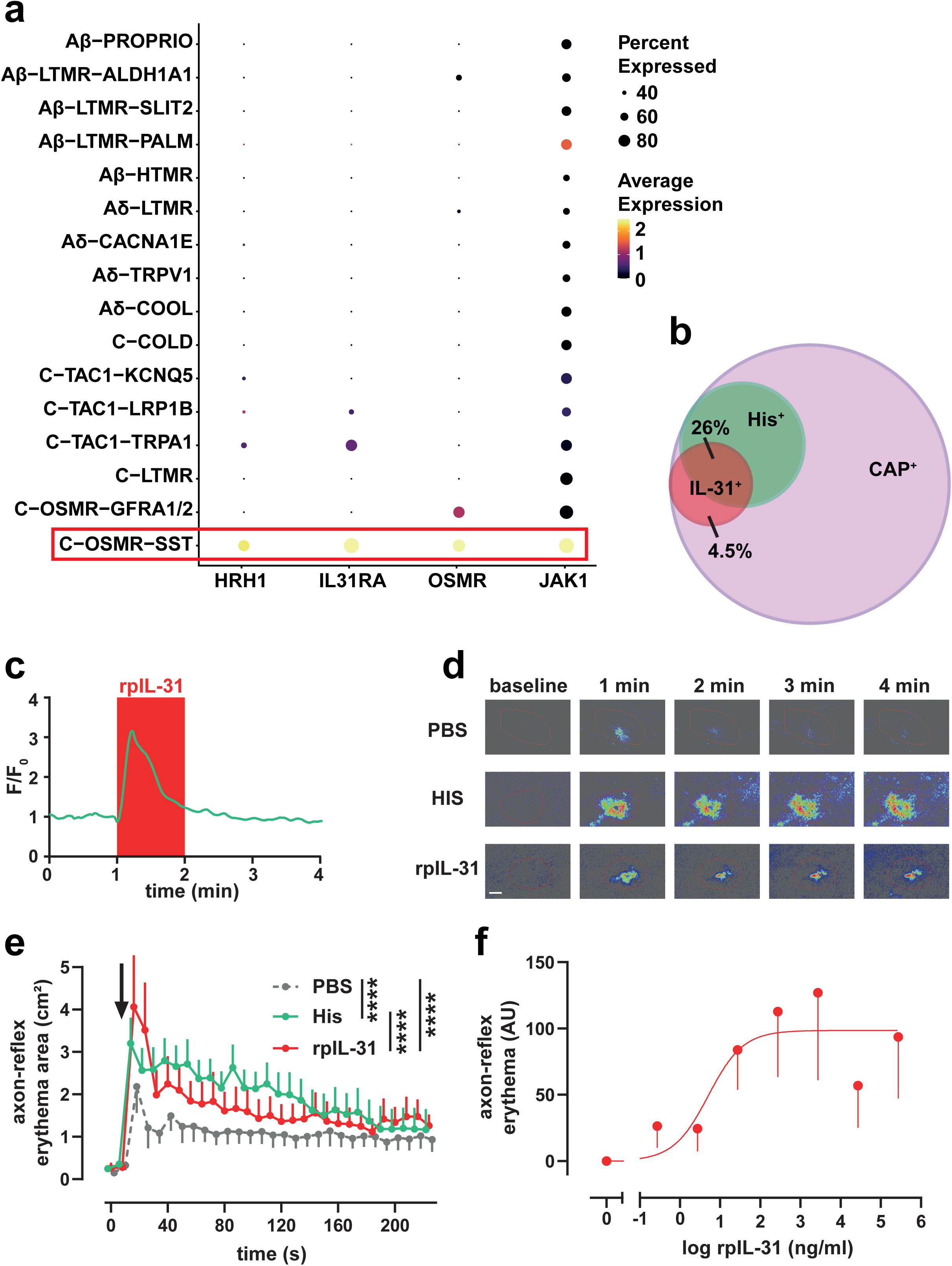
A small proportion of porcine DRG neurons is directly activated by rpIL31. a) Expression of *HRH1*, *IL31RA*, *OSMR* and *JAK1* in pig DRG neuron types, as assessed by reanalysis of snRNAseq data (13)of porcine thoracic and lumbar DRGs shown by a dot plot. C-OSMR-SST neurons co-expressing high levels of *HRH1* and the IL-31 receptor complex are highlighted with a red rectangle. b) Venn diagram showing the proportion of rpIL-31-responsive neurons inside the CAP^+^/His^+^ and CAP^+^ neuron types (n = 3 pigs). Note that rpIL-31 responses were not detected in CAP^−^ neurons. In total, we measured rpIL-31 responses in 23 CAP⁺/His⁺, 85 CAP⁺ and 94 CAP⁻ neurons. c) Representative calcium trace showing a fast-onset strong rpIL-31 response in a CAP^+^/His^+^ neuron. The rpIL-31 stimulation period is visualized by red shading. Note that the maximum calcium level during rpIL-31 stimulation was reached ∼15 seconds post stimulation onset. d) Laser speckle real-time flux images depict perfusion changes in pig skin in 1-min intervals at baseline and 4 minutes post injection of PBS, histamine (10 mg/ml) and rpIL-31 (272 ng/ml). Image sequences are representative for injections in 5 pigs (9-10 total injections per group). Scale bar = 1 cm. e) Axon-reflex erythema area (mean + SEM) plotted against time for injections with PBS, histamine and rpIL-31 (labelled with a black arrow), repeated measures mixed-effect analysis with Sidak’s post-hoc test. f) Integrated axon-reflex erythema (mean – SEM, see methods for calculation) is plotted as a function of injected rpIL-31 doses (n = 4-10). Based on a 3-parameter sigmoidal fit, we estimate an apparent EC_50_ value of 5 ± 10 ng/ml.

### Sensitization of porcine DRG neurons by IL-31

Based on IL-31 receptor complex expression patterns, many more CAP^+^/His^+^ neurons should be able to respond to rpIL-31 than those producing an immediate calcium transient (Fig. 4). Keeping with the established role of IL-31 in chronic itch and high intradermal IL-31 levels in chronic itch patients, we hypothesized that IL-31 potentially sensitizes porcine DRG neurons such that a subsequent chemical or electrical stimulation would result in stronger activation. Therefore, we adapted a protocol used by Tseng et al. in mouse DRG neurons to test whether rpIL-31 stimulation modifies responsiveness to repetitive histamine treatment (19). In detail, we stimulated porcine DRG neurons twice with histamine, 4 minutes apart, and recorded the induced calcium transients. Under control conditions, when we washed the cells between both histamine stimulation periods with imaging buffer, we noticed a substantial decrease in response amplitude to the second stimulation (Fig. 5A, left). However, when the cells were treated for 1 minute with rpIL-31 in between both histamine stimulation periods, this tachyphylaxis was largely abrogated (Fig. 5A, right). The response to the second histamine stimulation, relative to the first, was only reduced to 69 ± 5% after rpIL-31, while this value was reduced to 45 ± 9% during control conditions (Fig. 5B; p = 0.0245, t-test). Likewise, the percentage of responding cells was only reduced to 72% after rpIL-31 compared to 22% under control conditions (Fig. 5C; p = 0.0369, Fisher’s exact test). Of note, the reduction in histamine tachyphylaxis was observed in the absence of an acute rpIL-31-induced calcium transient (Fig. 5A), indicating that the underlying signaling pathway is indeed independent of intracellular calcium mobilization. Collectively, acute treatment with rpIL-31 robustly reduced the tachyphylaxis to repeated histamine stimulation in CAP^+^/His^+^ neurons. Under chronic conditions, this pathway might contribute to chronic itch, if additional pruritogens are affected in a similar fashion.

**Figure 5.**
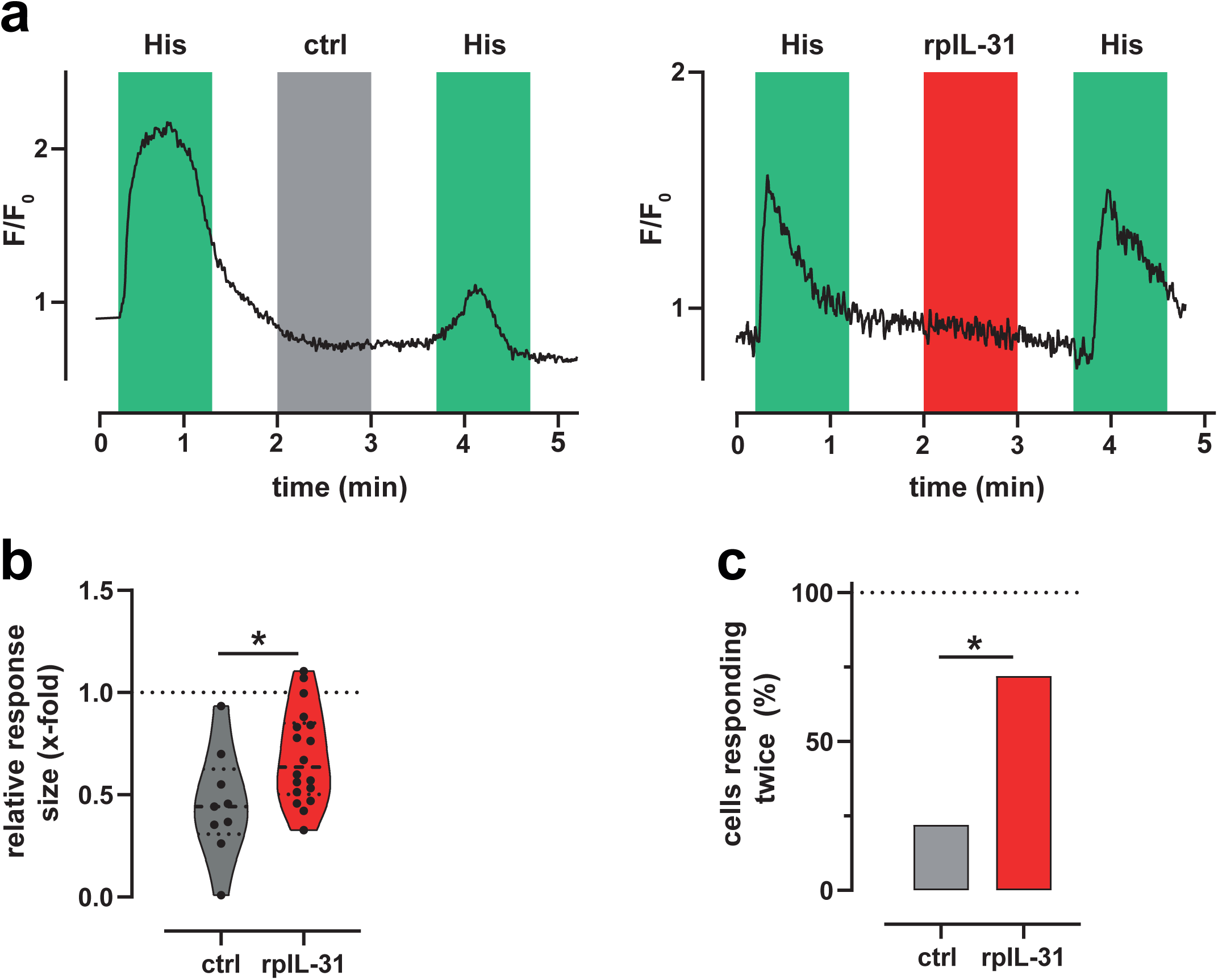
rpIL-31 reduces tachyphylaxis to repetitive histamine stimulation in porcine CAP^+^/His^+^ DRG neurons. a) Calcium traces show responses to two consecutive histamine applications (200 µM, green shaded area) in exemplary CAP^+^/His^+^ neurons. Intermitted, imaging buffer as control (left, grey shaded area) or rpIL-31 (272 ng/ml, right, red shaded area) was applied. Data is representative for n = 9 (ctrl) and n = 18 (rpIL-31) neurons from 6 pigs. b) Calcium transient response sizes to the second histamine application relative to the response sizes to the first histamine application under control conditions (grey) or after intermitted rpIL-31 treatment (red) is depicted as violin plot with dots representing single neurons (dashed line: median, dotted lines: quartiles), unpaired two-sided t-test. c) Proportion of neurons responding to the second histamine application relative to the number of neurons responding to the first histamine application under control conditions (grey) or after intermitted rpIL-31 treatment (red), Fisher’s exact test.

As a more general test for a potential rpIL-31-induced neuronal sensitization, we studied, whether the same short treatment with rpIL-31 increases electrical excitability. First, we repeated stimulation with 1, 2 and 4 rectangular pulses (100 mA) twice with and without intermitted treatment with rpIL-31 (Fig. S3A+B). For statistical comparisons we included neurons responding to all three stimulations during the first stimulation sequence and calculated relative changes of the net size of the induced calcium transients to the second stimulation sequence. We observed a reduction of the response size to repeated rectangular electrical stimulation already under control conditions that did not show neuron-type specificity for any number of rectangular pulses (Fig. S3C-E, H (8) = 12.90, p = 0.1153, Kruskal-Wallis). Importantly, rpIL-31 did neither enhance or reduce the size (all p > 0.1070, Kruskal-Wallis) of the second calcium transient in any neuron group for any number of stimulation pulses. Collectively, we therefore conclude that calcium transients induced by high intensity rectangular electrical stimulation of porcine DRG neurons are largely unaffected by acute rpIL-31 treatment.

Second, we also studied effects of rpIL-31 on excitability to sinusoidal electrical stimulation. In detail, we measured thresholds to slow and fast sine waves before and after rpIL-31 treatment (Fig. 6A+B). We first compared changes in response sizes of calcium transients induced by both sine waves at high stimulation intensity (50 mA), including all neurons responding during the first stimulation sequence to the respective sine wave frequency (Fig. S4A+B). However, we did not find rpIL-31-induced changes in net response sizes (all p > 0.5292, Kruskal-Wallis with Dunn’s post-hoc test) in any neuron group to high intensity stimulation with both sine waves. This is in good agreement with rectangular electrical stimulation results and further emphasizes that calcium signals induced by a low number of high intensity electrical stimuli of any wave form are largely unaffected by rpIL-31 in porcine DRG neurons. Therefore, we also analyzed whether rpIL-31 treatment sensitized sine wave thresholds. Including all cells with a detectable threshold to either of the two sine waves during the first stimulation sequence, we noticed that more than 25% of all CAP^+^/His^+^ neurons acquired detectable thresholds to fast sine in addition to slow sine wave stimulation after rpIL-31 treatment, while these neurons only had measurable thresholds to slow sine wave stimulation before. To quantify this effect, we substituted the missing threshold values by an arbitrary threshold (55 mA) slightly above our maximum stimulus intensity (50 mA) and calculated threshold reductions by rpIL-31 and under control conditions. Indeed, in CAP^+^/His^+^ neurons rpIL-31 induced a clear threshold reduction in comparison to controls after fast sine but not after slow sine wave stimulation, where thresholds stayed essentially the same (Fig. 6C+D). This behavior reached statistical significance (fast sine: p = 0.0279; slow sine: p = 0.8198, Mann-Whitney U) and was not detected in the other two neuron types (Fig. S4C). However, because effect size estimation is problematic with censored outcomes (42, 43) as observed in our case with cells lacking a threshold during the first stimulation sequence, we also employed Bayesian censored regression to more appropriately estimate rpIL-31 effects in our porcine DRG neuron groups. Our results essentially corroborated the findings from simple substitutions of missing threshold values (Fig. 6E+F). We found that rpIL-31 reduced the fast but not the slow sine wave threshold in a neuron-type dependent manner, as only CAP^+^/His^+^ neurons showed a threshold reduction by ∼10 mA with a posterior probability of 0.94. Collectively, this indicated that, in addition to enhancing repetitive chemical activation with pruritogens, rpIL-31 can also increase electrical excitability in CAP^+^/His^+^ neurons. However, we detected the latter effect only at threshold levels for a non-preferred wave form of this neuronal class.

**Figure 6.**
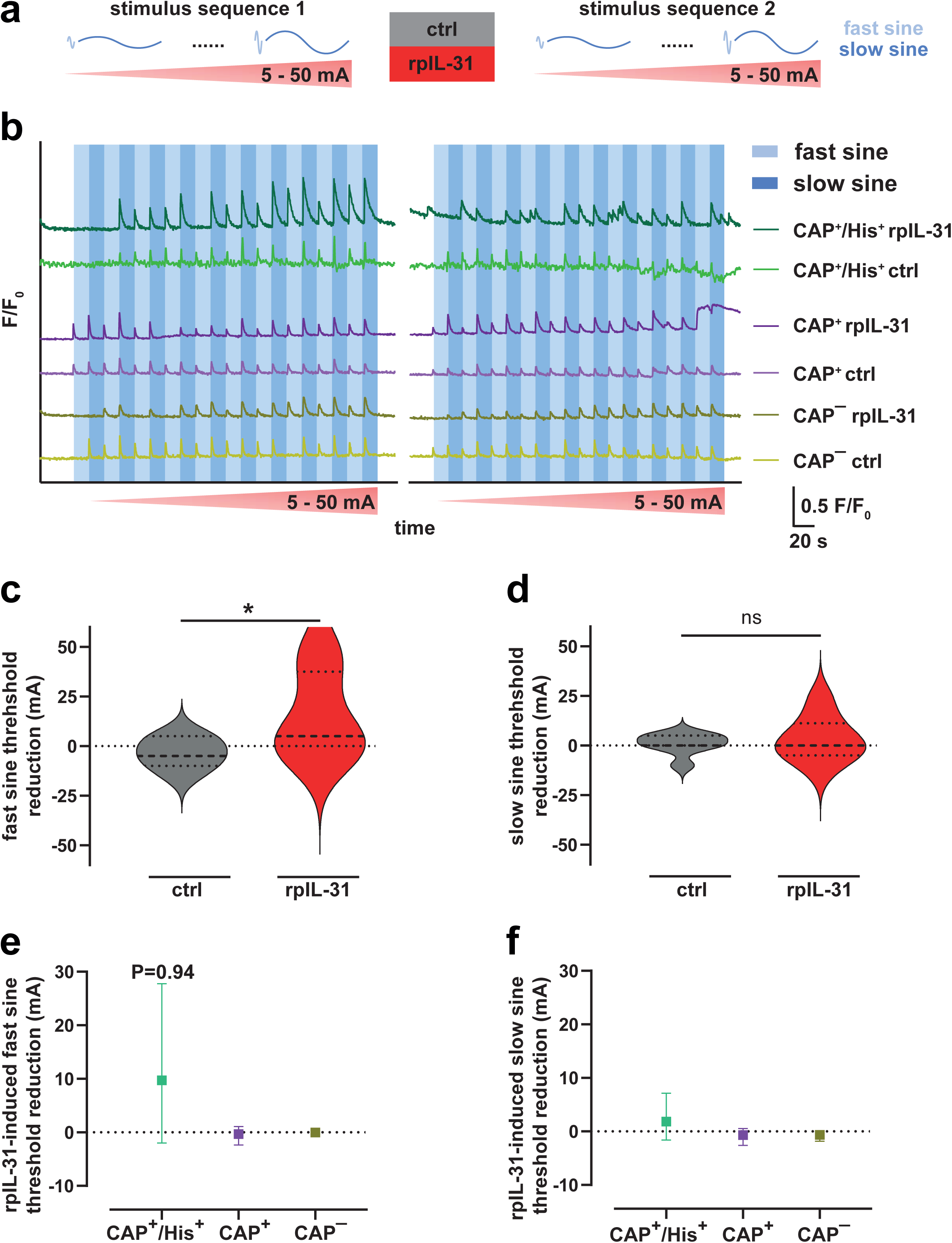
Modulation of calcium transients elicited by sinusoidal electrical stimulation in porcine DRG neurons by rpIL-31. a) Sinusoidal electrical stimulation protocol consisting of a repeated stimulation sequence with increasing stimulation intensity from 5 to 50 mA in 5 mA steps, alternating between slow and fast sine waves at each intensity level, without (ctrl) and with intermitted rpIL-31 (272 ng/ml, n = 6 pigs). b) Calcium traces from individual neurons representative for the three neuron types show responses to sinusoidal electrical stimulation before and after rpIL-31 application or under control conditions. The timing of sinusoidal electrical stimulation (5 - 50 mA in 5 mA steps, alternating between slow and fast sine waves at each intensity level) is indicated by lighter and darker blue shading, respectively. c) Reduction of the threshold to repeated fast sinusoidal electrical stimulation (20 ms) under control conditions (grey) or after intermitted rpIL-31 treatment (red) in CAP^+^/His^+^ neurons. Data for cells with a threshold to fast and/or slow sine wave stimulation during the first stimulation sequence (n = 24) was included. Missing threshold values in non-responders were set to 55 mA. Data is depicted as violin plot (dashed line: median, dotted lines: quartiles), Mann-Whitney U test. d) Reduction of the threshold to repeated slow sinusoidal electrical stimulation (250 ms) under control conditions (grey) or after intermitted rpIL-31 treatment (red) for same CAP^+^/His^+^ neurons shown in C. Missing threshold values in non-responders were set to 55 mA. Data is depicted as violin plot (dashed line: median, dotted lines: quartiles), Mann-Whitney U test. e) Reduction of the threshold to repeated fast sinusoidal electrical stimulation under control conditions or after intermitted rpIL-31 treatment was estimated with a Bayesian censored mixed-effects regression model in neurons with a threshold to fast and/or slow sine wave stimulation during the first stimulation sequence (n = 24 (CAP^+^/His^+^), n = 57 (CAP^+^), and n = 151 (CAP^−^)). The difference in threshold reduction (rpIL-31 – ctrl) is depicted as mean ± credible interval from 8000 model simulations. Note the high posterior probability of a ∼10 mA threshold reduction in CAP^+^/His^+^ neurons (P = 0.94). e) Reduction of the threshold to repeated slow sinusoidal electrical stimulation under control conditions or after intermitted rpIL-31 treatment was estimated with a Bayesian censored mixed-effects regression model in the same neurons included in E. The difference in threshold reduction (rpIL-31 – ctrl) is depicted as mean ± credible interval from 8000 model simulations.

## Discussion

The successful classification of primary afferent nerve fiber classes based on single-cell expression patterns has provided a new perspective to sensory physiology as such classifications were originally based purely on unique response characteristics. Currently, there is a major task to link the functional and transcriptomically defined classifications. Our data links the cluster of neurons characterized by selective expression of *SST*, *HRH1*, *OSMR, IL31RA* and *NPPB* to mechano-insensitive histamine-responsive pruriceptors based not only on chemical response patterns but coherently also on characteristic excitability patterns originally established to identify such fibers *in vivo*.

### Expression pattern vs. functional classification

In an initial classic approach, we used calcium responses to agonists of selectively expressed receptors to functionally classify cultured DRG neurons. Among those, CAP was used to target TRPV1+ nociceptive neurons. While in the mouse there is a major proportion of TRPV1^−^ nociceptors, in humans most of the nociceptors express *TRPV1* (44–46). Accordingly, local injections of CAP into human skin leave an “analgesic bleb” (47) indicating a minor role or even absence of TRPV1^−^ nociceptors. In broad agreement, with the exception of potential cold nociceptors, *TRPV1* transcripts were also detected in all transcriptionally defined neuron clusters putatively annotated with nociceptive function in a recent single-cell RNAseq study of porcine DRG neurons (13).Thus, for human and pig, CAP responses can be assumed to identify nociceptive primary afferent neurons. In addition, we used histamine to preferentially target neurons with high *HRH1* expression levels and a prime role in histaminergic itch. Such a link may appear problematic based on non-specific histamine responses. In humans, histamine iontophoresis strongly and selectively activates mechano-insensitive C-chemonociceptors, producing histamine itch (20). However, histamine injection also activates polymodal nociceptors, albeit only transiently (27), and such an activation of polymodal nociceptors can be linked to a transient and lower intensity burning pain after histamine injections (48). Although we cannot exclude that our histamine stimulus also activated other neuron types, assays with a high degree of signal amplification such as *in vitro* calcium imaging are generally well suited to separate cell types based on receptor expression and downstream signaling (49) and it is thus worth noting that C-OSMR-SST neurons harbor by far the highest *HRH1* expression level. About 90% of histamine responsive neurons in our study also responded to CAP being in good agreement with the RNAseq data for C-OSMR-SST neurons but this, unfortunately, cannot differentiate mechano-insensitive chemonociceptors from polymodal nociceptors.

Clinically, histamine-induced itch is particularly linked to acute itch upon mast cell degranulation such as during allergic reactions or in the context of urticaria, but beyond this, anti-histamines have no central role in chronic itch conditions such as atopic dermatitis (AD) or prurigo nodularis (50). One might therefore be tempted to conclude that also histamine-responsive pruriceptors are of minor importance in these conditions. Based on the clinical success of nemolizumab as antipruritic treatment in chronic dermatological itch (6, 7), the mechanism of IL-31-induced itch, including neuronal IL-31 responses, has been scrutinized with some positive results for acute activation (10, 15, 16), and major longer lasting effects on neuronal outgrowth (51) and sensitization (10, 18, 19). Importantly, the IL-31 receptor complex is a conserved marker of a particular neuronal cluster in human and pig, i.e. mechano-insensitive nociceptors(13), and such neurons also mediate histamine-induced itch in human skin (20, 52). Thus, while histamine does not play a key role in chronic pruritus of AD patients, histamine-responsive mechano-insensitive pruriceptors that are uniquely equipped with the IL-31 receptor complex are prime candidates to explain a direct anti-pruritic effect of nemolizumab. Our data functionally confirm the combined expression of histamine- and IL-31 receptor complex comprised of IL31RA and OSMRβ in a subpopulation of CAP^+^ neurons. While direct activation of these neurons by IL-31 was rare, it robustly reduced histamine tachyphylaxis. Thus, histamine responsive neurons were found co-responsive to IL-31, supporting the combined receptor expression in these neurons as predicted by single-cell transcriptomics only for the “itch cluster”.

### Basic electrophysiology

Extracellular electric field stimulation enabled functional classification of isolated porcine DRG neurons using the same paradigms as established to classify sensory afferents *in vivo* (*26*). Overall, we found that several of those *in vivo* characteristics appear to be maintained at “soma-level”. First, CAP^−^ non-nociceptors appear to be less prone to accumulation of calcium by trains of action potentials than CAP^+^ nociceptors. This feature is highly congruent with the shorter duration of action potentials, in part a consequence of the lower expression of NaV1.8, which harbors quite slow gating kinetics, and with the extremely high following frequencies of Aβ-fibers of ∼400 Hz (25, 53). However, the most striking findings of our study are differences among the CAP^+^ nociceptors separated by a positive histamine response. These, presumably pruriceptive neurons had much longer calcium transients (T50) associated with electrically elicited APs as compared to presumably non-pruriceptive nociceptors. This might impact on several calcium-dependent processes of pruriceptive neurons and we hypothesize that it is of particular importance for localized neuro-immune cross-talk, enabling e.g. calcium-dependent neuropeptide release during neurogenic inflammation (10, 54, 55). Beyond the classic combination of vasodilation and protein extravasation, a much more specific efferent function of C-OSMR-SST neurons can be expected for the release of brain natriuretic peptide (BNP, encoded by the *NPPB* gene) as it has been shown to induce a migratory phenotype in Langerhans cells (which reduce levels of the BNP receptor NPR1 in AD) (56), and is linked to itch and inflammation in mouse models of AD (57). Importantly, bi-directional neuroimmune interactions are expected, as increased pruriceptor-specific expression of *Nppb* has been shown in chronic itch models in mouse and monkey (58).

In addition, CAP^+^/His^+^ neurons also showed a strong preference for slow depolarizing electrical pulses (250 ms) as compared to fast 20 ms sine waves. Interestingly, this feature was unique for CAP^+^/His^+^ neurons, whereas no frequency preference was found in histamine negative nociceptors and non-nociceptors. This is remarkable as silent nociceptors, in comparison to classic polymodal nociceptors have similar thresholds to slow sinusoidal stimulation but much higher thresholds to fast rectangular depolarizing pulses *in vivo* in humans and pigs (23, 30, 59). In combination with recent patch-seq data (13), this congruent functional characteristic further supports the linkage of transcriptionally defined histamine positive “itch neurons” and a silent nociceptor-like behavior.

### Limitations

In our study, we measured calcium signals as a proxy for neuronal AP activity and not APs itself. Although we cannot exclude that this approach misrepresents neuronal activity, others have formally validated extracellular electric field stimulation of DRG neurons to elicit APs (60). In addition, in our study the step-like increase of calcium signals to increasing numbers of rectangular electrical stimuli as well as IL-31-induced calcium signals *in vitro* and axon-reflex erythema *in vivo* also support the validity of our approach. In the future, voltage imaging will provide a more nuanced approach to extend our current approach also to the sub-threshold voltage range. In addition, we focused on direct, short-term effects of IL-31. While this approach allowed drawing conclusions on immediate and direct IL-31 effects in sensory neurons, clinical translation will likely be influenced by multiple IL-31 receptor complex expressing non-neuronal cell types and engagement of long-term signal transduction pathways, including gene expression changes, by IL-31 in patient’s skin (2–5, 61).

### Translational implications

Our data suggest that neuronal IL-31 responses are found specifically in a particular class of mechano-insensitive and histamine-responsive pruriceptors. Spontaneous activity in this class of C-chemonociceptors has already been found in chronic itch and pain conditions (62–64). The combination of sparse direct activation by IL-31 and profound reduction of tachyphylaxis to repetitive pruritogen-stimulation of these neurons in culture appear to match the early anti-pruritic onset of nemolizumab in AD and prurigo patients even before the reduction of the inflammatory eczema (6, 7). While this anti-pruritic effect is observed on a time scale of hours, IL-31-induced sprouting of sensory nerve endings is assumed to contribute with a tardy time-course of days (51), but has not been investigated in our study. It is important to note that the crucial role of counteracting neuronal sensitization by anti-pruritic therapy is also reflected in the recently revised definition of chronic itch, that acknowledges the change of peripheral and central itch processing (65).

In summary, our data reflects a successful back-translational approach from humans to pigs that considered species differences, such as lack of cross-species activity of IL-31, to functionally characterize a particular neuronal class of pruriceptors. To enable mechanistic studies relying on molecular genetics, it is an important future challenge to also back-translate these results to rodents, where transcriptional similarity in general (31) and co-expression of *Il31ra*, *Osmr*, *Sst*, *Hrh1*, and *Nppb* as marker genes in particular (66–68) is conserved in a single neuron type originally coined NP3, yet the functional equivalent, dermal mechano-insensitive C-chemonociceptors are not present in the skin (69–71). Given this challenge, direct neuron-mediated effects of other anti-pruritic drugs, e.g. Janus kinase inhibitors (72, 73), might prospectively also be studied in pigs, harnessing their high degree of both, transcriptional and functional similarity to humans.

## Materials and methods

### Tissue extraction

Pig dorsal root ganglia (DRG) were collected according to the 3R criteria to minimize animal use as surplus tissue from previously completed independent experiments (ethical clearance obtained from *Regierungspräsidium Karlsruhe* under the protocol numbers G-78/18 and G-99/23). To this aim, 12 male domestic pigs (*Sus scrofa domesticus*) with an average age of 12 weeks and a body weight of 20-25 kg, were euthanized in deep pentobarbital (30 mg/kg body weight) anaesthesia by intravenous administration of a supersaturated potassium chloride solution (1 ml/kg). Six to nine DRGs from thoracic and lumbar segments were dissected and placed in ice-cold HBSS (with Ca^2+^/Mg^2+^; ThermoFisher, Karlsruhe, Germany).

### Pig DRG neuron culture

Glass cover slips (25 mm round, Carl Roth, Germany) were placed in 35 mm diameter culture dishes (Thermo Fisher) and pre-coated for 30-90 minutes with Laminin (5 µg/ml; Sigma Aldrich, St. Louis, MO, USA) and poly-L-lysine (0.1%, molecular weight 70.000-150.000 Da; Sigma Aldrich). Next, a small droplet of diluted Laminin (150 µg/ml in sterile water) was applied to the centre of each cover slip and allowed to air-dry such that the resulting adhesive area for subsequent cell seeding stayed visible due to the formation of a white precipitate. Harvested DRGs in HBSS were cleaned by trimming connected roots and nerves. Depending on the size of the ganglion, each DRG was cut into 3–6 pieces, and the fragments were then enzymatically digested in Liberase™ DH (250 µg/ml in HBSS; Roche, Mannheim, Germany) for 40 minutes at 37 °C. Afterwards, 27.5 µL of DNase I (7.4 mg/ml, Sigma) was added, and the tube was returned to 37 °C for an additional 5 minutes. Following the removal of 500 µl of supernatant, 1 ml of Trypsin-EDTA (0.25%, ThermoFisher) was added and incubated for 7 minutes at 37 °C. Enzymatic activity was stopped by adding 1 ml of complete medium (CM), consisting of DMEM/F-12 supplemented with 10% fetal bovine serum (FBS), 100 U/ml penicillin and 100 µg/ml streptomycin (all Life Technologies, Darmstadt, Germany). Cells were gently triturated with a 1 ml micropipette tip until a cloudy suspension was obtained and filtered through a 300 µm cell strainer (pluriSelect, Leipzig, Germany). The filtrate was centrifuged (2 min, 260 x *g*, room temperature), and after resuspension of the pellet in CM, cells were separated from myelin debris via a 15.5% Percoll (Sigma Aldrich) cushion (5 min, 720 x *g*, room temperature). Cells present in the pellet phase, were washed twice by resuspension in CM and subsequent centrifugation for 2 min at 260 x *g* and room temperature. Before seeding, cells were resuspended in medium containing 20% FBS and approximately 30 µl of the cell suspension was seeded directly onto the laminin-coated white area of each cover slip. Cells were allowed to attach for 1.5–2 h at 37 °C/5% CO_2_ before each culture dish was flushed with 2 ml of CM supplemented with 100 ng/ml rhβNGF. Cells were kept at 37 °C/5% CO_2_ overnight for measurement on the following day.

### Live cell calcium imaging

#### Solutions

The tests were conducted using an imaging buffer containing 140 mM NaCl, 4 mM KCl, 2 mM CaCl_2_, 2 mM MgCl_2_, 10 mM HEPES, and 15 mM D-glucose (pH 7.4, adjusted with NaOH). Stimulation solutions were prepared by diluting stock solutions in imaging buffer to the following final concentrations: 200 µM histamine dihydrochloride (Sigma Aldrich), 272 ng/ml rpIL-31 (GenScript, Piscataway NJ, USA), 10 µM capsaicin (Sigma Aldrich), and high-K⁺ solution (replacing 55 mM NaCl in the imaging buffer with 55 mM KCl). Histamine (10 mM in imaging buffer), Capsaicin (100 mM in DMSO) and rpIL-31 (272 µg/ml in PBS) stock solutions were stored in aliquots at –20 °C, 4 °C and -80 °C, respectively.

#### Imaging setup

Calcium imaging was performed as previously described (74). Briefly, cells were loaded with the calcium tracer Fluo-8AM (3 μM, AAT Bioquest, Sunnyvale CA, USA) in imaging buffer for 20-40 minutes at room temperature protected from light. Afterwards, cover slips were mounted in a slotted bath (RC-21 BRFS, Warner Instruments, Holliston MA, USA) equipped with two platinum wire electrodes to allow extracellular electrical stimulation. Subsequently, the chamber was placed on an inverted Axiovert 200 (Zeiss, Oberkochen, Germany) epifluorescence microscope equipped with a custom-built gravity-fed 7-channel perfusion system for chemical stimulation. Fluorescence signals (exposure time 10-12 ms) were acquired at 4 Hz with an Evolve 512 camera (Photometrics, Tucson AZ, USA) using a 465 nm LED (Prior Scientific Instruments, Jena, Germany) as excitation light source, a 510 nm dichroic mirror and a 515 nm long-pass emission filter. Illumination and image acquisition were controlled by Micro-Manager software.

#### Electrical stimulation

Bipolar constant current electrical field stimulation was delivered by an isolated constant current stimulator (DS5, Digitimer, UK) that was connected to a pulse generator (NI USB-6221, National Instruments, USA). DAPSYS8.0 software was used to control electrical stimulation, allowing rectangular (amplitude 100 mA, 4 Hz) and complete sine (amplitude 5-50 mA, wavelength 250 and 20 ms) wave pulses delivered via the two electrodes of the stage-mounted slotted bath. If not otherwise indicated, only a single stimulus was given.

#### Data analysis

Image sequences were analysed with ImageJ software (version 1.54fj; (75)). Circular regions of interest (ROIs) corresponding to individual neuronal somata were manually drawn. Only cells that responded to capsaicin or high-K^+^ solution, applied at the end of each experiment, were included in the analysis. The average background fluorescence, calculated for each time-point from five cell-free background ROIs distributed over the whole field of view, was subtracted from the fluorescence intensity (F) measured in all cell-based ROIs. For each cell ROI, the fluorescence values were subsequently normalized to the mean baseline fluorescence (F_0_) of the first 20 frames (F/F_0_).

The response of a cell to electrical stimulation was determined by a two-factor thresholding approach. First, fluorescence during stimulation (8-10 frames) was compared to the fluorescence just before the stimulation (10 frames) using a one-tailed t-test and a response was accepted at p < 0.05. Second, a cell’s enhanced fluorescence during stimulation had to pass a ΔF/F_0_ threshold of 0.05. Cells that had met both criteria were labelled as responsive. Chemical responses were considered positive when exceeding the pre-decided threshold criterion of average baseline fluorescence before the stimulation plus 5 standard deviations (20 frames). Cells with spontaneous calcium transients during the baseline period were filtered out.

For statistical group comparisons we calculated net response sizes (ΔF/F_0_) and duration (T50) of induced calcium transients. In detail, ΔF/F_0_ is the maximum fluorescence during the stimulation period minus the average fluorescence of the prior baseline period. T50 is the decay time from the maximum net increase to 50% of that maximum signal. Slow and fast sine wave thresholds were determined by identifying the lowest stimulation amplitude intensity that produced a positive response according to the two-factor thresholding approach detailed above. The slow sine wave preference of a cell was calculated by subtracting the threshold of slow from the threshold of fast sine wave stimulation (Thsh_fast_ – Thsh_slow_).

### snRNAseq data reanalysis

A barcode-gene expression-matrix with fully annotated neuronal barcodes was acquired from the gene expression omnibus under accession number GSE263532 (13). Dot plots to depict per cell-type gene expression for the following genes were created in R (76) using the packages Seurat (77) and scCustomize (78): *HRH1*, *TRPV1*, *IL31RA*, *OSMR*, and *JAK1*.

### Skin perfusion measurements

According to the 3R criteria, male domestic pigs (age 12 weeks) under pentobarbital anaesthesia were used for non-invasive skin perfusion measurements after independent experiments were completed (ethical clearance obtained from *Regierungspräsidium Karlsruhe* under the protocol number G-99/23). A laser speckle contrast imager (FLPI-2, Moor instruments, UK) was positioned ∼50 cm over the intended measurement area such that the scanning light path was perpendicular to the skin surface. Measurements were conducted in 5 pigs using both, rump and upper forelimb skin. Imaging in temporal mode was controlled via the Moor FLPI imaging software with the following settings: sample rate 0.13 Hz, exposure time 20 ms, time constant 1.0 s, temporal filter 100 frames. Baseline perfusion was assessed for 30 seconds before 10 µl of PBS as negative control (n = 10), histamine as positive control (10 mg/ml, n = 10) or various doses of rpIL-31 (272 pg/ml to 27.2 µg/ml in 10-fold dilution steps, n = 4-10) were injected intradermally using a 31G insulin syringe. Imaging was continued for a total of 5 minutes. To determine the area of enhanced skin perfusion for each image, a thresholding approach inside a hand-drawn region of interest was used as previously described (79). The resulting axon-reflex erythema area was plotted against time. For overall statistical comparison, we selected the most effective rpIL-31 dose (272 ng/ml) and compared groups with a repeated measures mixed-effect model with time and treatment as fixed effects. This analysis indicated a highly significant treatment effect (F (2, 459) = 86.17, p < 0.0001), while time played no significant role (F (26, 243) = 1.356, p = 0.1227) and interaction of treatment and time (F (52, 459) = 1.531, p = 0.0129) had only a minor effect. To establish a dose-response relationship, rpIL-31-induced enhanced skin perfusion was calculated for all rpIL-31 doses by integrating the axon-reflex erythema area vs time curve from 0.5 to 4 minutes, subtracting the per animal negative control value and plotting this specific rpIL-31 effect against the rpIL-31 dose. To estimate an EC_50_ dose, this dataset was fit with a 3-parameter sigmoidal curve (GraphPad Prism 8.4.3: log(agonist) vs. response (three parameter)), constraining the bottom to 0 due to the negative control subtraction.

### Reporter gene assay

#### Plasmids

pGL4.47 (Promega, Walldorf, Germany) was used to measure activity of the STAT3 transcription factor. It encodes a hPEST-destabilized version of firefly (*Photinus pyralis*) luciferase under a minimal promoter and five copies of the STAT3 response element SIE. This configuration allows a sensitive detection of STAT3-activating signalling pathways after relatively short stimulation periods. pGL4.74 (Promega) encodes an octocorallia (*Renilla reniformis*) luciferase under the general HSV-TK promoter. pcDNA3.1-pigIL31RA-T2A-pigOSMR-T2A-TurboRFP encodes a single open reading frame for a fusion protein of porcine IL31RA (UniProt accession F1SLL2), porcine OSMRβ (UniProt accession F1SN77) and TurboRFP. The addition of the self-cleavage peptide T2A between the three proteins allows the generation of separate functional proteins. This plasmid was custom synthetized and sequence-verified by GenScript (Piscataway, USA).

#### Cell culture and transfection

HEK293 cells were cultured in DMEM high glucose medium supplemented with 10% FBS, 2 mM L-glutamine, penicillin and streptomycin (all Life Technologies). One day before transfection, 100,000 cells were seeded into 12-well dishes. Cells were transfected with Promofectin (Promocell, Germany) according to the manufacturer’s instructions, using per well a mixture of 100 ng pGL4.74, 1 µg pGL4.47 with and without 1 µg pcDNA3.1-pigIL31RA-T2A-pigOSMR-T2A-TurboRFP.

#### Dual luciferase assay

1 day post transfection, cells were serum-starved in DMEM high glucose medium without FBS for 16 hours. Then, cells were stimulated in duplicate in 1 ml DMEM high glucose medium with or without various concentrations of rpIL-31 or with 250 ng/ml recombinant human IL-31 (rhIL-31, ThermoFisher). After 6 hours stimulation, cell culture medium was removed and cells were washed once with 1 ml PBS (Life Technologies). Cell lysis was performed in 100 µl of 1X passive lysis buffer (Promega) by rocking for 15 minutes at room temperature. 50 µl lysate was transferred to a white bottom 96-well plate (PerkinElmer, Rodgau, Germany) and the plate was placed in a FluoStar Omega luminescence reader (BMG Labtech, Ortenberg, Germany). First, total firefly luciferase luminescence was measured at 2 Hz for 20 seconds by injection of 50 µl luciferase assay reagent II (Promega) with a computer-controlled pump of the luminescence reader, 1 second after start of the measurement. Second, total *Renilla* luciferase luminescence was measured, again at 2 Hz for 20 seconds, by injection of 50 µl Stop & Glo reagent (Promega) with a second computer-controlled pump of the luminescence reader, 1 second after start of the second measurement. Luminescence values for both enzymes were averaged during the 3-20 second period of each measurement and STAT3 reporter activity was calculated as the ratio of firefly by *Renilla* luminescence. To depict changes in STAT3 reporter activity, activity in treated cells was normalized to non-treated cells, separately for each transfection condition. To establish an EC_50_ dose for rpIL-31-induced STAT3 reporter activation, data from separate experiments was normalized and plotted against the rpIL-31 dose, by first fitting with a 4-parameter sigmoidal curve (GraphPad Prism 8.4.3: log(agonist) vs. response – variable slope (four parameter)), constraining the bottom to 1 and then expressing the reporter activity as % of the fitted top plateau.

### Statistics

For all *in vitro* experiments, individual neurons (ROIs) were considered as biological replicates. For *in vivo* experiments, individual intradermal injections were considered as biological replicates. Exact replicate numbers as well as the number of animals used are given in the respective figure legend. Group data is presented in violin plots with median and quartiles indicated by dotted lines or as mean with SEM to indicate variance, as indicated in figure legends. Calcium traces of individual neurons representative for the analysed groups and exemplary skin flux images are presented.

Normal distribution of data was tested with a Shapiro-Wilk test and by visually assessing QQ plots. Accordingly, when data did not follow normal distribution group comparisons of calcium transient response sizes, T50 values and slow sine wave preference were conducted with non-parametric Kruskal-Wallis ANOVA and pair-wise Dunn’s post-hoc tests, when > 2 groups were compared, and experiments harboring only 2 groups were compared by two-sided t-tests. Population sizes of responding/non-responding neurons were compared across groups using chi-square tests or within the neuron group using Fisher’s exact or chi-square tests for trend. *In vivo* skin perfusion data was analyzed with a repeated-measures mixed-effect model and pair-wise Sidak’s post-hoc test. Due to missing thresholds during sine wave electrical stimulation, a Bayesian censored (Tobit-type) mixed-effects regression model was fit using the R package brms (version 2.23.0) as this has been shown to be superior to simple substitutions (42) and is highly relevant for threshold change estimations under treatment (43). Censored (missing) thresholds were assumed to underly a latent continuous lognormal distribution. Fixed effects included stimulus repetition, neuron group and rpIL-31 treatment, repetition x neuron group and rpIL-31 treatment x neuron group as interactions. In addition, a cell level random effect was included. Bayesian inference used Hamiltonian Monte Carlo simulations with weak-informative priors and the following arguments of the brm()-function: init = 0, chains = 4, iter = 8000, warmup = 2000, adapt_delta = 0.9999, max_treedepth = 15. For all statistical comparisons, significant differences were accepted at pre-defined p-values and are depicted in figures using asterisks (*: p < 0.05, **: p < 0.01, ***: p < 0.001, ****: p < 0.0001). In addition, exact p-values are reported when appropriate in the text. If not otherwise indicated, GraphPad Prism 8 (Dotmatics, Boston MA, USA) was used for data plotting and for statistical analysis.

## Ethics Statement

Ethical clearances for animal experiments presented in this manuscript were obtained from *Regierungspräsidium Karlsruhe* under the protocol number G-99/23.

## Data Availability Statement

The authors affirm that all data necessary for confirming the conclusions of the article are present within the article, figures, and tables.

## Acknowledgments

The authors thank Elmar Forsch for expert technical assistance with DRG extraction and skin perfusion measurements. The authors gratefully acknowledge funding support by DFG, German Research Foundation through grant 255156212/CRC1158/Z02-INF and 350193106/FOR2690/TP01 to H.J.S., through grant 255156212/CRC1158/A01 and 350193106/FOR2690/TP01 to M.S. and through grant 350193106/FOR2690/TP03 and 397846571 to R.R.. Funding sources were not involved in study design, data collection, analysis and interpretation, manuscript writing and the decision to submit the manuscript for publication.

## Author Contributions Statement

Conceptualization: Z.A., M.S., H.J.S.

Data Curation: Z.A., H.J.S.

Formal Analysis: Z.A., H.J.S., M.S.

Funding Acquisition: M.S., H.J.S.

Investigation: Z.A., H.J.S.

Methodology: M.B.

Project Administration: M.S., H.J.S.

Resources: S.S., R.R.

Supervision: M.S., H.J.S.

Validation: M.S., H.J.S.

Visualization: Z.A., H.J.S.

Writing – Original Draft Preparation: Z.A., M.S., H.J.S.

Writing – Review and Editing: Z.A., M.B., S.S., R.R., M.S., H.J.S.

## Abbreviations

DRG: dorsal root ganglion
ANOVA: analysis of variances
AP: action potential
AD: atopic dermatitis
CM: complete medium
ROI: region of interest

## Tables

**Supplemental Table 1.** Evolutionary similarity of IL-31 on the protein level*.

|  | **Human** | **Pig** | **Mouse** |
| --- | --- | --- | --- |
| **Human** |  | 56.9 | 48.0 |
| **Pig** | 56.9 |  | 45.4 |
| **Mouse** | 48.0 | 45.4 |  |

\* Evolutionary conservation of IL-31 was assessed by pairwise sequence alignment using the Needleman-Wunsch algorithm for global alignment with standard parameters. Similarity scores (%) were calculated using the BLOSUM62 substitution matrix. Primary amino acid sequences were downloaded from UniProt using the accession numbers Q6EBC2 (human), F1RNT4 (pig) and Q6EAL8 (mouse).

**Supplemental Table 2.** Evolutionary similarity of IL31RA on the protein level*.

|  | **Human** | **Pig** | **Mouse** |
| --- | --- | --- | --- |
| **Human** |  | 82.5 | 71.1 |
| **Pig** | 82.5 |  | 68.3 |
| **Mouse** | 71.1 | 68.3 |  |

\* Evolutionary conservation of IL31RA was assessed by pairwise sequence alignment using the Needleman-Wunsch algorithm for global alignment with standard parameters. Similarity scores (%) were calculated using the BLOSUM62 substitution matrix. Primary amino acid sequences were downloaded from UniProt using the accession numbers Q8NI17 (human), F1SLL2 (pig) and Q8K5B1 (mouse).

**Supplemental Table 3.** Evolutionary similarity of OSMRβ on the protein level*.

|  | **Human** | **Pig** | **Mouse** |
| --- | --- | --- | --- |
| **Human** |  | 82.5 | 69.3 |
| **Pig** | 82.5 |  | 68.8 |
| **Mouse** | 69.3 | 68.8 |  |

\* Evolutionary conservation of OSMRβ was assessed by pairwise sequence alignment using the Needleman-Wunsch algorithm for global alignment with standard parameters. Similarity scores (%) were calculated using the BLOSUM62 substitution matrix. Primary amino acid sequences were downloaded from UniProt using the accession numbers Q99650 (human), F1SN77 (pig) and O70458 (mouse).

## Supplemental figure legends

**Figure S1** AP-associated calcium transients increase less with increasing stimulation pulse number in CAP^−^ neurons.

Calcium transient response sizes associated with 1/2/4 pulses of 100 mA rectangular electrical stimulation applied at 4 Hz (raw data shown in Figure 2C) were normalized on a per-cell level to responses to a single stimulation pulse and are depicted as violin plot (dashed line: median, dotted lines: quartiles). Data was analysed with ANOVA (F (5, 354) = 11.02, p < 0.0001) and pair-wise comparisons with Sidak’s post-hoc test within each pulse number group. We detected a significant difference only between CAP^+^ and CAP^−^ neurons, when stimulated with 4 pulses (p = 0.0005).

**Figure S2** Recombinant pig IL-31 activates pig IL-31 receptor complex in a heterologous expression system.

a) STAT3 reporter activity was measured in HEK293 cells with or without (Mock) overexpression of porcine IL31RA and OSMRβ (n = 3) and is depicted as violin plot (dashed line: median, dotted lines: quartiles) on a log_2_-scale. Cells were stimulated in duplicate with rpIL-31 (270 ng/ml) or recombinant human IL-31 (rhIL-31, 250 ng/ml) or left unstimulated (ctrl). Data was analysed with ANOVA (F (4, 10) = 39.82, p < 0.0001) and pairwise comparisons with a Sidak’s post-hoc test indicated that rpIL-31 significantly stimulated STAT3 reporter activity only if its cognate receptor complex was expressed (Mock: p > 0.9999; + pig IL31RA/OSMRβ: p < 0.0001). rhIL-31 did not cross-activate pig IL31RA/OSMRβ (p > 0.9999).

b) STAT3 reporter activity was measured in HEK293 cells overexpressing porcine IL31RA and OSMRβ (n = 4). Cells were stimulated in duplicate with various concentrations of rpIL-31.

Normalized STAT3 reporter activity (mean ± SEM) is plotted against the rpIL-31 concentration. Based on a 4-parameter sigmoidal fit, we estimate an apparent EC_50_ value of 48 ± 5 ng/ml.

**Figure S3** Modulation of AP-associated calcium transients in porcine DRG neurons by rpIL-31.

a) Electrical stimulation protocol consisting of a repeated stimulation sequence with 1, 2 and 4 rectangular pulses, each without (ctrl) and with intermitted rpIL-31 (272 ng/ml, n = 6 pigs).

b) Calcium traces from individual neurons representative for the three neuron types show responses to rectangular electrical stimulation before and after rpIL-31 application or under control conditions. The timing of rectangular electrical pulses (100 mA) is indicated by grey shading with increasing intensity, from light (1 pulse) over medium (2 pluses) to dark grey (4 pulses). Data is representative for n = 19 (CAP^+^/His^+^), n = 46 (CAP^+^) and n = 89 (CAP^−^) neurons.

c-e) Calcium transient response sizes to a repeated sequence of rectangular electrical stimulation with 1, 2 and 4 pulses relative to a first stimulation sequence with similar properties under control conditions (grey) or after intermitted rpIL-31 treatment (red). Data was split by neuron group (c, CAP^+^/His^+^ neurons; d, CAP^+^ neurons; e, CAP^−^ neurons) and is presented as violin plots (dashed line: median, dotted lines: quartiles), Kruskal-Wallis ANOVA.

**Figure S4** Calcium response sizes elicited by strong sinusoidal electrical stimulation in all porcine DRG neuron types and thresholds to fast sinusoidal electrical stimulation in CAP^+^ and CAP^−^ neurons are not changed by rpIL-31.

a) Calcium transient response sizes to repeated fast sinusoidal electrical stimulation (20 ms) with high intensity (50 mA), relative to an initial stimulation with similar properties, under control conditions (grey) or after intermitted rpIL-31 treatment (272 ng/ml, red). Data for cells responding during the first stimulation sequence (n = 11 (CAP^+^/His^+^), n = 42 (CAP^+^), and n = 97 (CAP^−^)) was included and is depicted as violin plot (dashed line: median, dotted lines: quartiles), Kruskal-Wallis ANOVA with Dunn’s post-hoc test.

b) Calcium transient response sizes to repeated slow sinusoidal electrical stimulation (250 ms) with high intensity (50 mA), relative to an initial stimulation with similar properties, under control conditions (grey) or after intermitted rpIL-31 treatment (272 ng/ml, red). Data for cells responding during the first stimulation sequence (n = 22 (CAP^+^/His^+^), n = 48 (CAP^+^), and n = 119 (CAP^+^)) was included and is depicted as violin plot (dashed line: median, dotted lines: quartiles), Kruskal-Wallis ANOVA with Dunn’s post-hoc test.

c) Reduction of the threshold to repeated fast sinusoidal electrical stimulation under control conditions (grey) or after intermitted rpIL-31 treatment (red). Missing threshold values in non-responders were set to 55 mA. Data for cells with a threshold to fast and/or slow sinusoidal stimulation during the first stimulation sequence (n = 24 (CAP^+^/His^+^), n = 57 (CAP^+^), and n = 151 (CAP^−^)) was included and is depicted as violin plot (dashed line: median, dotted lines: quartiles), Kruskal-Wallis ANOVA.

## References

1. Coscarella G, Edwards E, Yosipovitch G. Basic Mechanisms of Itch and Advances in Clinical Management. Ann Allergy Asthma Immunol. 2025.

2. Hashimoto T, Nattkemper LA, Kim HS, Kursewicz CD, Fowler E, Shah SM, et al. Itch intensity in prurigo nodularis is closely related to dermal interleukin-31, oncostatin M, IL-31 receptor alpha and oncostatin M receptor beta. Experimental dermatology. 2021;30(6):804–10.

3. Kim S, Kim HJ, Yang HS, Kim E, Huh IS, Yang JM. IL-31 Serum Protein and Tissue mRNA Levels in Patients with Atopic Dermatitis. Ann Dermatol. 2011;23(4):468–73.

4. Kremer AE, Mayo MJ, Hirschfield GM, Levy C, Bowlus CL, Jones DE, et al. Seladelpar treatment reduces IL-31 and pruritus in patients with primary biliary cholangitis. Hepatology. 2024;80(1):27–37.

5. Singer EM, Shin DB, Nattkemper LA, Benoit BM, Klein RS, Didigu CA, et al. IL-31 is produced by the malignant T-cell population in cutaneous T-Cell lymphoma and correlates with CTCL pruritus. J Invest Dermatol. 2013;133(12):2783–5.

6. Kwatra SG, Yosipovitch G, Legat FJ, Reich A, Paul C, Simon D, et al. Phase 3 Trial of Nemolizumab in Patients with Prurigo Nodularis. The New England journal of medicine. 2023;389(17):1579–89.

7. Silverberg JI, Wollenberg A, Reich A, Thaci D, Legat FJ, Papp KA, et al. Nemolizumab with concomitant topical therapy in adolescents and adults with moderate-to-severe atopic dermatitis (ARCADIA 1 and ARCADIA 2): results from two replicate, double-blind, randomised controlled phase 3 trials. Lancet. 2024;404(10451):445–60.

8. Sofen H, Bissonnette R, Yosipovitch G, Silverberg JI, Tyring S, Loo WJ, et al. Efficacy and safety of vixarelimab, a human monoclonal oncostatin M receptor beta antibody, in moderate-to-severe prurigo nodularis: a randomised, double-blind, placebo-controlled, phase 2a study. EClinicalMedicine. 2023;57:101826.

9. Solinski HJ, Dranchak P, Oliphant E, Gu X, Earnest TW, Braisted J, et al. Inhibition of natriuretic peptide receptor 1 reduces itch in mice. Sci Transl Med. 2019;11(500).

10. Meng J, Moriyama M, Feld M, Buddenkotte J, Buhl T, Szollosi A, et al. New mechanism underlying IL-31-induced atopic dermatitis. J Allergy Clin Immunol. 2018;141(5):1677–89 e8.

11. Jung M, Dourado M, Maksymetz J, Jacobson A, Laufer BI, Baca M, et al. Cross-species transcriptomic atlas of dorsal root ganglia reveals species-specific programs for sensory function. Nat Commun. 2023;14(1):366.

12. Cevikbas F, Wang X, Akiyama T, Kempkes C, Savinko T, Antal A, et al. A sensory neuron-expressed IL-31 receptor mediates T helper cell-dependent itch: Involvement of TRPV1 and TRPA1. J Allergy Clin Immunol. 2014;133(2):448–60.

13. Korner J, Howard D, Solinski HJ, Mancilla Moreno M, Haag N, Fiebig A, et al. Molecular architecture of human dermal sleeping nociceptors. Cell. 2026.

14. Hawro T, Saluja R, Weller K, Altrichter S, Metz M, Maurer M. Interleukin-31 does not induce immediate itch in atopic dermatitis patients and healthy controls after skin challenge. Allergy. 2014;69(1):113–7.

15. Lewis KE, Holdren MS, Maurer MF, Underwood S, Meengs B, Julien SH, et al. Interleukin (IL) 31 induces in cynomolgus monkeys a rapid and intense itch response that can be inhibited by an IL-31 neutralizing antibody. J Eur Acad Dermatol Venereol. 2017;31(1):142–50.

16. Oyama S, Kitamura H, Kuramochi T, Higuchi Y, Matsushita H, Suzuki T, et al. Cynomolgus monkey model of interleukin-31-induced scratching depicts blockade of human interleukin-31 receptor A by a humanized monoclonal antibody. Experimental dermatology. 2018;27(1):14–21.

17. Pearson J, Leon R, Starr H, Kim SJ, Fogle JE, Banovic F. Establishment of an Intradermal Canine IL-31-Induced Pruritus Model to Evaluate Therapeutic Candidates in Atopic Dermatitis. Vet Sci. 2023;10(5).

18. Takahashi S, Ochiai S, Jin J, Takahashi N, Toshima S, Ishigame H, et al. Sensory neuronal STAT3 is critical for IL-31 receptor expression and inflammatory itch. Cell Rep. 2023;42(12):113433.

19. Tseng PY, Hoon MA. Oncostatin M can sensitize sensory neurons in inflammatory pruritus. Sci Transl Med. 2021;13(619):eabe3037.

20. Schmelz M, Schmidt R, Bickel A, Handwerker HO, Torebjork HE. Specific C-receptors for itch in human skin. J Neurosci. 1997;17(20):8003–8.

21. Fiebig A, Leibl V, Oostendorf D, Lukaschek S, Frombgen J, Masoudi M, et al. Peripheral signaling pathways contributing to non-histaminergic itch in humans. J Transl Med. 2023;21(1):908.

22. Ringkamp M, Schepers RJ, Shimada SG, Johanek LM, Hartke TV, Borzan J, et al. A role for nociceptive, myelinated nerve fibers in itch sensation. J Neurosci. 2011;31(42):14841–9.

23. Jonas R, Namer B, Stockinger L, Chisholm K, Schnakenberg M, Landmann G, et al. Tuning in C-nociceptors to reveal mechanisms in chronic neuropathic pain. Annals of neurology. 2018;83(5):945–57.

24. Werland F, Hirth M, Rukwied R, Ringkamp M, Turnquist B, Jorum E, et al. Maximum axonal following frequency separates classes of cutaneous unmyelinated nociceptors in the pig. The Journal of physiology. 2020.

25. Johansson RS, Landstrom U, Lundstrom R. Responses of mechanoreceptive afferent units in the glabrous skin of the human hand to sinusoidal skin displacements. Brain Res. 1982;244(1):17–25.

26. Rukwied R, Thomas C, Obreja O, Werland F, Kleggetveit IP, Jorum E, et al. Slow depolarizing stimuli differentially activate mechanosensitive and silent C nociceptors in human and pig skin. Pain. 2020;161(9):2119–28.

27. Namer B, Carr R, Johanek LM, Schmelz M, Handwerker HO, Ringkamp M. Separate peripheral pathways for pruritus in man. Journal of neurophysiology. 2008;100(4):2062–9.

28. Obreja O, Ringkamp M, Namer B, Forsch E, Klusch A, Rukwied R, et al. Patterns of activity-dependent conduction velocity changes differentiate classes of unmyelinated mechano-insensitive afferents including cold nociceptors, in pig and in human. Pain. 2010;148(1):59–69.

29. Schneider T, Filip J, Soares S, Sohns K, Carr R, Rukwied R, et al. Optimized Electrical Stimulation of C-Nociceptors in Humans Based on the Chronaxie of Porcine C-Fibers. The journal of pain : official journal of the American Pain Society. 2023;24(6):957–69.

30. Dileep D, Bali K, Soares S, Schmelz M, Rukwied R. Optimum electrical sinusoidal frequency stimulation to activate C-nociceptors. Pain Rep. 2026;11(2):e1412.

31. Bhuiyan SA, Xu M, Yang L, Semizoglou E, Bhatia P, Pantaleo KI, et al. Harmonized cross-species cell atlases of trigeminal and dorsal root ganglia. Sci Adv. 2024;10(25):eadj9173.

32. Nguyen MQ, von Buchholtz LJ, Reker AN, Ryba NJ, Davidson S. Single-nucleus transcriptomic analysis of human dorsal root ganglion neurons. Elife. 2021;10.

33. Yu H, Nagi SS, Usoskin D, Hu Y, Kupari J, Bouchatta O, et al. Leveraging deep single-soma RNA sequencing to explore the neural basis of human somatosensation. Nature neuroscience. 2024;27(12):2326–40.

34. Summerfield A, Meurens F, Ricklin ME. The immunology of the porcine skin and its value as a model for human skin. Mol Immunol. 2015;66(1):14–21.

35. Uhm C, Jeong H, Lee SH, Hwang JS, Lim KM, Nam KT. Comparison of structural characteristics and molecular markers of rabbit skin, pig skin, and reconstructed human epidermis for an ex vivo human skin model. Toxicol Res. 2023;39(3):477–84.

36. Kim YK, Lee JY, Hwang JH, Suh HN. A Pilot Study To Establish an Ovalbumin-induced Atopic Dermatitis Minipig Model. J Vet Res. 2021;65(3):307–13.

37. Vana G, Meingassner JG. Morphologic and immunohistochemical features of experimentally induced allergic contact dermatitis in Gottingen minipigs. Vet Pathol. 2000;37(6):565–80.

38. Meijs S, Schmelz M, Meilin S, Jensen W. A systematic review of porcine models in translational pain research. Lab Anim (NY). 2021;50(11):313–26.

39. Cornelissen C, Luscher-Firzlaff J, Baron JM, Luscher B. Signaling by IL-31 and functional consequences. Eur J Cell Biol. 2012;91(6-7):552–66.

40. Schmelz M, Michael K, Weidner C, Schmidt R, Torebjörk HE, Handwerker HO. Which nerve fibers mediate the axon reflex flare in human skin? Neuroreport. 2000;11(3):645–8.

41. Lynn B, Schutterle S, Pierau FK. The vasodilator component of neurogenic inflammation is caused by a special subclass of heat-sensitive nociceptors in the skin of the pig. JPhysiol(Lond). 1996;494(Pt 2):587–93.

42. Lubin JH, Colt JS, Camann D, Davis S, Cerhan JR, Severson RK, et al. Epidemiologic evaluation of measurement data in the presence of detection limits. Environ Health Perspect. 2004;112(17):1691–6.

43. Angst MS, Phillips NG, Drover DR, Tingle M, Ray A, Swan GE, et al. Pain sensitivity and opioid analgesia: a pharmacogenomic twin study. Pain. 2012;153(7):1397–409.

44. Rostock C, Schrenk-Siemens K, Pohle J, Siemens J. Human vs. Mouse Nociceptors - Similarities and Differences. Neuroscience. 2018;387:13–27.

45. Schmelz M, Schmid R, Handwerker HO, Torebjork HE. Encoding of burning pain from capsaicin-treated human skin in two categories of unmyelinated nerve fibres. Brain. 2000;123 Pt 3:560–71.

46. Shiers S, Klein RM, Price TJ. Quantitative differences in neuronal subpopulations between mouse and human dorsal root ganglia demonstrated with RNAscope in situ hybridization. Pain. 2020;161(10):2410–24.

47. LaMotte RH, Shain CN, Simone DA, Tsai EF. Neurogenic hyperalgesia: psychophysical studies of underlying mechanisms. Journal of neurophysiology. 1991;66(1):190–211.

48. Sikand P, Shimada SG, Green BG, LaMotte RH. Sensory responses to injection and punctate application of capsaicin and histamine to the skin. Pain. 2011;152(11):2485–94.

49. de Melo Reis RA, Freitas HR, de Mello FG. Cell Calcium Imaging as a Reliable Method to Study Neuron-Glial Circuits. Front Neurosci. 2020;14:569361.

50. Thurmond RL, Kazerouni K, Chaplan SR, Greenspan AJ. Antihistamines and itch. Handb Exp Pharmacol. 2015;226:257–90.

51. Feld M, Garcia R, Buddenkotte J, Katayama S, Lewis K, Muirhead G, et al. The pruritus- and TH2-associated cytokine IL-31 promotes growth of sensory nerves. J Allergy Clin Immunol. 2016;138(2):500–8 e24.

52. Schmelz M, Michael K, Weidner C, Schmidt R, Torebjork HE, Handwerker HO. Which nerve fibers mediate the axon reflex flare in human skin? Neuroreport. 2000;11(3):645–8.

53. Djouhri L, Fang X, Okuse K, Wood JN, Berry CM, Lawson SN. The TTX-resistant sodium channel Nav1.8 (SNS/PN3): expression and correlation with membrane properties in rat nociceptive primary afferent neurons. The Journal of physiology. 2003;550(Pt 3):739–52.

54. Feng X, Zhan H, Sokol CL. Sensory neuronal control of skin barrier immunity. Trends Immunol. 2024;45(5):371–80.

55. Saraiva-Santos T, Zaninelli TH, Pinho-Ribeiro FA. Modulation of host immunity by sensory neurons. Trends Immunol. 2024;45(5):381–96.

56. Horvath D, Penzes Z, Molnar P, Rebenku I, Vereb G, Szanto M, et al. Natriuretic peptides modulate monocyte-derived Langerhans cell differentiation and promote a migratory phenotype. Front Immunol. 2025;16:1593141.

57. Wheeler JJ, Williams N, Yu J, Mishra SK. Brain Natriuretic Peptide Exerts Inflammation and Peripheral Itch in a Mouse Model of Atopic Dermatitis. J Invest Dermatol. 2024;144(3):705–7.

58. Tseng PY, Liew HL, Ringkamp M, Nagao K, Hoon MA. Upregulated natriuretic peptide B expression as a hallmark of chronic itch. Pain. 2025;166(12):2818–30.

59. Schmidt R, Schmelz M, Weidner C, Handwerker HO, Torebjork HE. Innervation territories of mechano-insensitive C nociceptors in human skin. Journal of neurophysiology. 2002;88(4):1859–66.

60. Fouillet A, Watson JF, Piekarz AD, Huang X, Li B, Priest B, et al. Characterisation of Nav1.7 functional expression in rat dorsal root ganglia neurons by using an electrical field stimulation assay. Mol Pain. 2017;13:1744806917745179.

61. Nemmer JM, Kuchner M, Datsi A, Olah P, Julia V, Raap U, et al. Interleukin-31 Signaling Bridges the Gap Between Immune Cells, the Nervous System and Epithelial Tissues. Front Med (Lausanne). 2021;8:639097.

62. Kleggetveit IP, Namer B, Schmidt R, Helas T, Ruckel M, Orstavik K, et al. High spontaneous activity of C-nociceptors in painful polyneuropathy. Pain. 2012;153(10):2040–7.

63. Serra J, Bostock H, Sola R, Aleu J, Garcia E, Cokic B, et al. Microneurographic identification of spontaneous activity in C-nociceptors in neuropathic pain states in humans and rats. Pain. 2012;153(1):42–55.

64. Schmelz M, Hilliges M, Schmidt R, Orstavik K, Vahlquist C, Weidner C, et al. Active “itch fibers” in chronic pruritus. Neurology. 2003;61(4):564–6.

65. Stander S, Schmelz M, Lerner E, Murota H, Nattkemper L, Reich A, et al. A multidisciplinary Delphi consensus on the modern definition of pruritus: Sensation and disease. J Eur Acad Dermatol Venereol. 2025.

66. Mishra SK, Hoon MA. The cells and circuitry for itch responses in mice. Science (New York, NY. 2013;340(6135):968–71.

67. Solinski HJ, Kriegbaum MC, Tseng PY, Earnest TW, Gu X, Barik A, et al. Nppb Neurons Are Sensors of Mast Cell-Induced Itch. Cell Rep. 2019;26(13):3561–73 e4.

68. Usoskin D, Furlan A, Islam S, Abdo H, Lonnerberg P, Lou D, et al. Unbiased classification of sensory neuron types by large-scale single-cell RNA sequencing. Nature neuroscience. 2015;18(1):145–53.

69. Kress M, Koltzenburg M, Reeh PW, Handwerker HO. Responsiveness and functional attributes of electrically localized terminals of cutaneous C-fibers in vivo and in vitro. Journal of neurophysiology. 1992;68(2):581–95.

70. Prato V, Taberner FJ, Hockley JRF, Callejo G, Arcourt A, Tazir B, et al. Functional and Molecular Characterization of Mechanoinsensitive “Silent” Nociceptors. Cell Rep. 2017;21(11):3102–15.

71. Qi L, Iskols M, Shi D, Reddy P, Walker C, Lezgiyeva K, et al. A mouse DRG genetic toolkit reveals morphological and physiological diversity of somatosensory neuron subtypes. Cell. 2024;187(6):1508–26 e16.

72. Oetjen LK, Mack MR, Feng J, Whelan TM, Niu H, Guo CJ, et al. Sensory Neurons Co-opt Classical Immune Signaling Pathways to Mediate Chronic Itch. Cell. 2017;171(1):217–28 e13.

73. Vazquez E, Richter F, Natura G, Konig C, Eitner A, Schaible HG. Direct Effects of the Janus Kinase Inhibitor Baricitinib on Sensory Neurons. Int J Mol Sci. 2024;25(22).

74. Behrendt M, Solinski HJ, Schmelz M, Carr R. Bradykinin-Induced Sensitization of Transient Receptor Potential Channel Melastatin 3 Calcium Responses in Mouse Nociceptive Neurons. Front Cell Neurosci. 2022;16:843225.

75. Schindelin J, Arganda-Carreras I, Frise E, Kaynig V, Longair M, Pietzsch T, et al. Fiji: an open-source platform for biological-image analysis. Nature methods. 2012;9(7):676–82.

76. R Development Core Team. R: A language and environment for statistical computing. 2010.

77. Hao Y, Stuart T, Kowalski MH, Choudhary S, Hoffman P, Hartman A, et al. Dictionary learning for integrative, multimodal and scalable single-cell analysis. Nat Biotechnol. 2024;42(2):293–304.

78. Marsh SE. scCustomize: Custom Visualizations & Functions for Streamlined Analyses of Single Cell Sequencing. version 3.2.0 ed2021.

79. Rukwied R, Schley M, Forsch E, Obreja O, Dusch M, Schmelz M. Nerve growth factor-evoked nociceptor sensitization in pig skin in vivo. J Neurosci Res. 2010;88(9):2066–72.

