## Supplementary figures and images for "Direct sensitizing and activating effects of interleukin 31 are restricted to a single, functionally and transcriptionally classified porcine DRG neuron subtype"

### Supplemental Figure 1

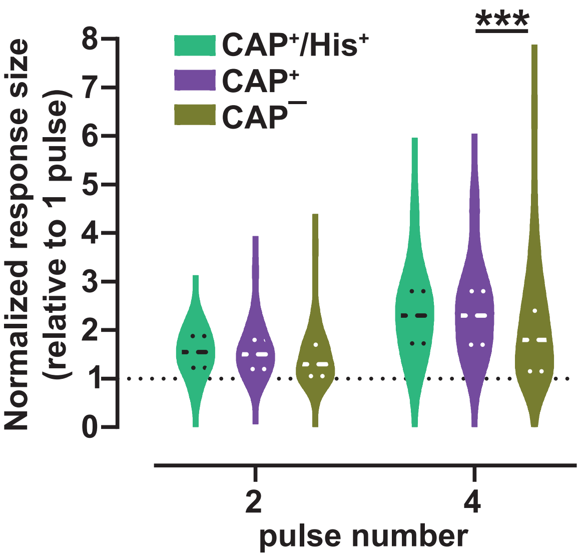

### Supplemental Figure 2

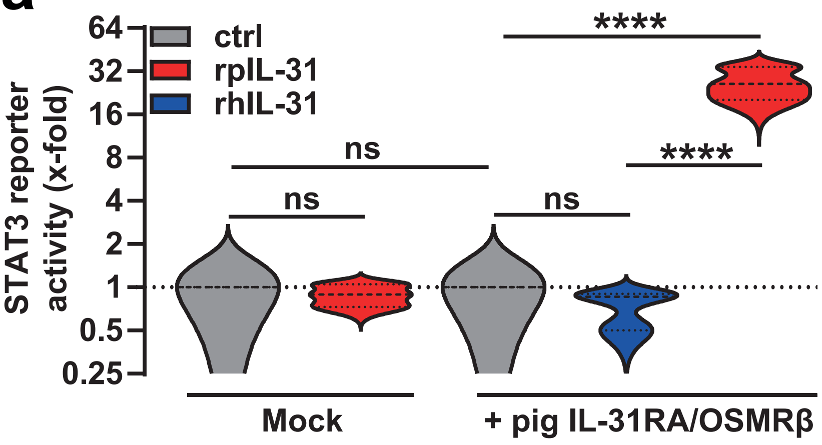

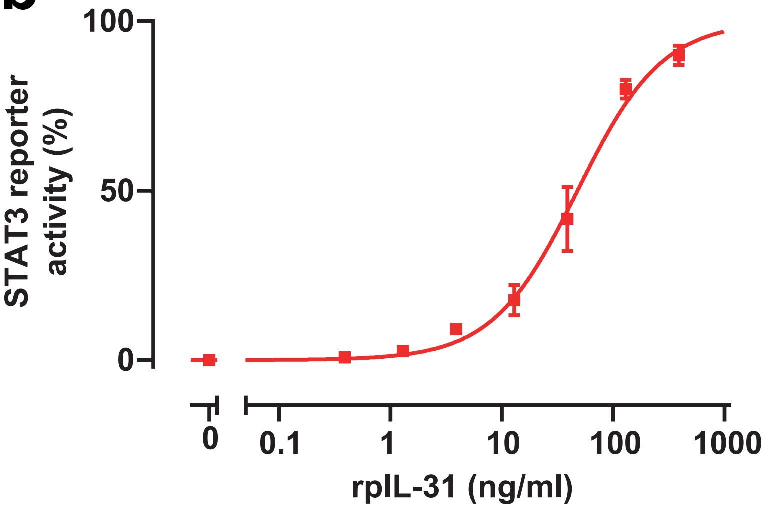

### Supplemental Figure 3

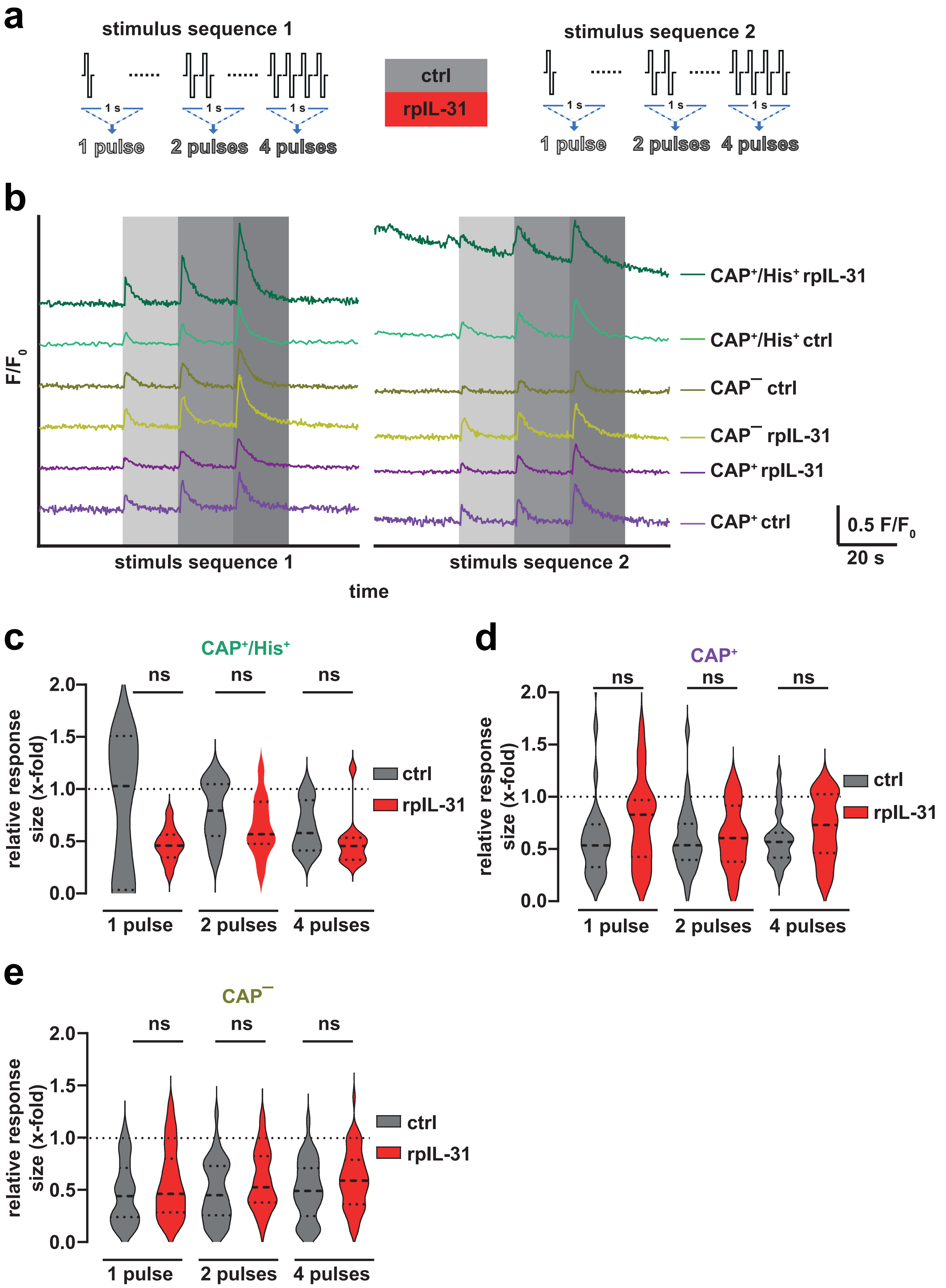

### Supplemental Figure 4

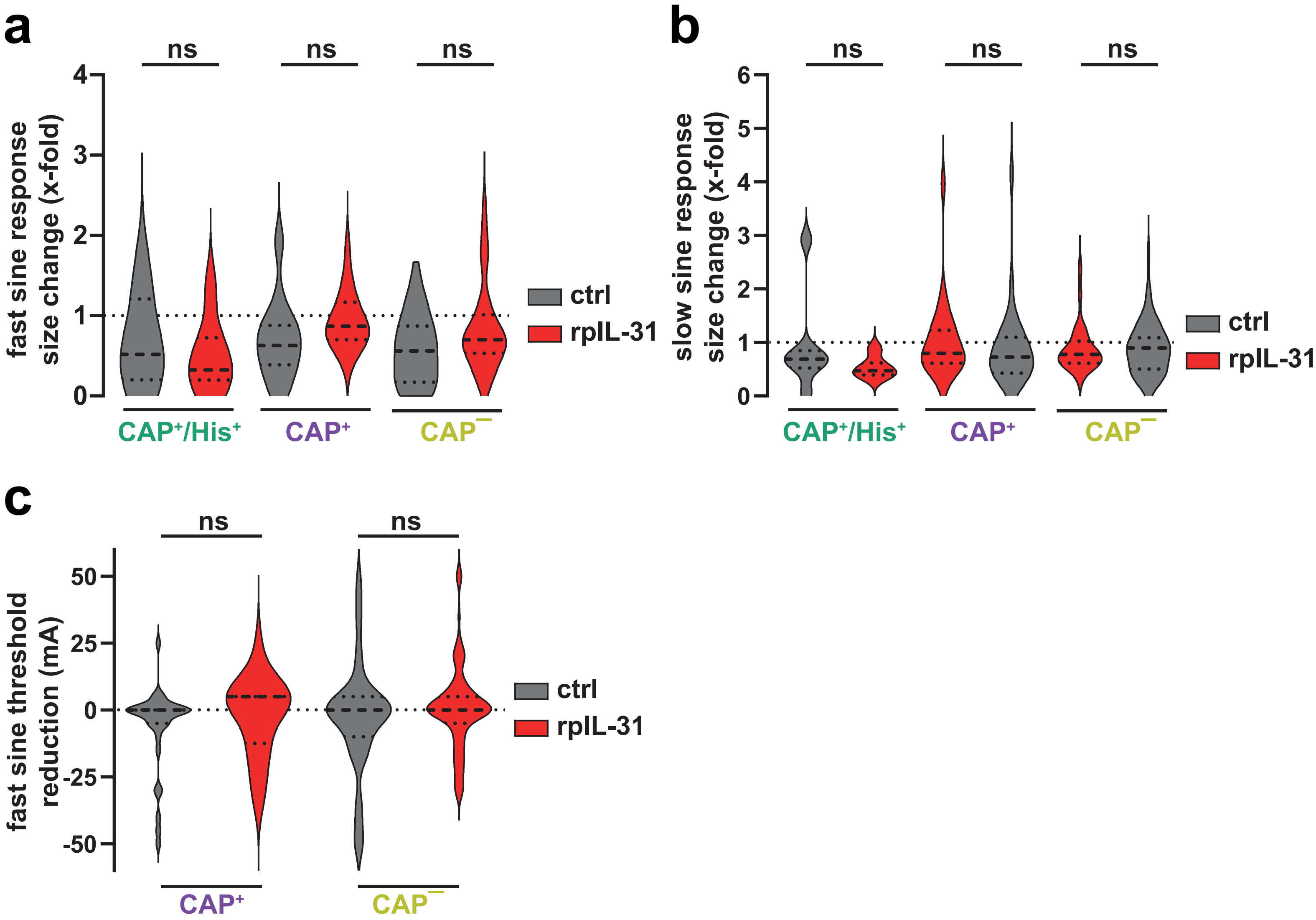
